# Among-population variation in the threespine stickleback (*Gasterosteus aculeatus*) liver metabolome: effects of ancestry and environment

**DOI:** 10.64898/2026.06.25.734390

**Authors:** Francis Quang-Hai Dinh, Mel Grillo, Marwa Mamdouh Tawfik, Andrew P. Hendry, Åsa Lind, Kathryn Milligan-McClellan, Catherine L. Peichel, Natalie C. Steinel, Yun-Jie Tseng, Jesse N. Weber, Monica Wu, Pieter C. Dorrestein, Andrés M. Caraballo-Rodríguez, Daniel I. Bolnick

## Abstract

Untargeted metabolomics offers a powerful lens for quantifying high-dimensional phenotypic variation within and among species in nature, but has yet to be widely adopted in evolutionary ecology. Some important initial questions are whether metabolome composition differs among populations, and to what extent such variation is genetic or plastic. Here, we use untargeted liquid chromatography tandem mass spectrometry to characterize the relative abundance of 5,939 molecular features of the threespine stickleback (*Gasterosteus aculeatus*) liver metabolome.

Native lake populations differ in metabolome composition, reflecting effects of sex, size, geography, and population ecotype (benthic versus limnetic). Stickleback from these lakes were translocated to found new populations in nine recently fishless lakes, permuting fish ecotypes across benthic and limnetic lake habitats. Several generations later, metabolomes in these experimental populations reflect effects both of their genetic ancestry (e.g., taurocholic acid, a cholane steroid bile acid, was elevated in limnetic-ancestries), as well as their present habitat (e.g., acylcarnitines). Additionally, ecotypes transplanted into a habitat to which they were maladapted exhibited a distinctive metabolomic profile. We conclude that stickleback exhibit both heritable and plastic among-population differences in liver metabolome, which could represent an important phenotypic basis of rapid evolution, population divergence, and perhaps local adaptation.

## INTRODUCTION

Metabolomics, or the comprehensive profiling of the small molecules within cells, tissues, and organisms, is well-suited for characterizing many aspects of an organism’s physiological phenotype (Oliver et al. 1998; Chen et al. 2025). These small molecules, known as metabolites, can be endogenously produced by genome-coded enzymatic processes, or exogenously obtained through diet, microbiota, and other environmental sources (Clish 2015). Consequently, the metabolome integrates genetic, epigenetic, and environmental influences into a single high-dimensional molecular readout of an organisms’ molecular phenotype (Fiehn 2002; Patti et al. 2012). Liquid-chromatography-tandem mass spectrometry (LC-MS/MS) is increasingly being used for ‘untargeted’ metabolomics, characterizing a broad profile of thousands of small chemicals within a given tissue. The approach is analogous to ‘shotgun’ sequencing for transcriptomics, in that it identifies most small molecules within a sample and estimates their relative abundance (Alanazi 2025). It is arguably a better measure of phenotypic traits than other -omics methods (Ryan & Robards 2017). Recent advances in instrumentation and computational tools have improved LC-MS/MS, enabling cost-effective and high-throughput molecular phenotyping (Alanazi 2025; Heuckeroth et al. 2024; Sarkar et al. 2024).

Despite rapid adoption of metabolomics in other disciplines, untargeted LC-MS/MS has not been widely adopted by evolutionary ecologists studying non-model organisms in nature. Untargeted LC-MS/MS metabolomics has the potential to provide quantitative phenotypic data on many traits relevant to evolution, adaptation, and ecological interactions, including immune responses (Wu et al. 2024), diet and nutrition (Marcias et al. 2024), temperature physiology (Xu et al. 2022), symbiosis (Coler et al. 2024, Hong et al. 2023), and stress. Metabolomic variation can be heritable, and thus evolvable, as revealed by genetic mapping studies in humans, poultry, and agricultural crops (Tahir et al. 2022; Tian et al. 2023; Li et al. 2023). To the extent that the metabolome responds plastically to environmental change, it can also provide insight into stress responses, acclimation, pollution, and reaction norm evolution (Lankadurai et al. 2013; Jeffries et al. 2016; Giebułtowicz et al. 2024). Yet, very few studies have used untargeted LC-MS/MS to document among-population variation in metabolomes in wild organisms (Ramanathan et al. 2026), or responses to environmental change (Bin et al. 2025), migration (Domer et al. 2024), and local adaptation (Wang et al. 2025). Although there is a small and growing literature comparing metabolomes across species in nature (Liu et al. 2021; Sparagon et al. 2022; Elsers et al. 2023; Wolters et al. 2026), we still lack rather basic information about intraspecific variation in metabolomes in nature. Does the metabolome differ among populations, within a species? Is metabolomic variation correlated with local environments, a pattern often used to infer adaptive function? To what extent are between-population differences in the metabolome environmentally induced, versus heritable (and thus evolvable)?

Here, we address these questions using untargeted LC-MS/MS data from 16 lake populations of threespine stickleback (*Gasterosteus aculeatus*), a popular evolutionary model organism (Bell and Foster, 1994, Reid et al. 2021; Strickland et al. 2021). Parallel evolution from a marine ancestor into freshwater-adapted populations, and divergence among freshwater populations, has produced extensive genetic and phenotypic variation (Jones et al. 2012; Stuart et al. 2017; Brachman et al. 2026). For example, lake stickleback using shallow littoral (benthic) versus open-water (limnetic) habitats exhibit characteristic differences in diet, morphology, behavior, physiology, and genotype. These differences are most pronounced in a few lakes containing benthic and limnetic species pairs (McPhail 1984; Schluter and McPhail, 1992), but many single-species lake populations also vary along this benthic-limnetic continuum (Lavin & MacPhail, 1985; Lavin & MacPhail, 1986; Matthews et al. 2010; Willacker et al. 2010; Bolnick and Ballare, 2020). This axis of variation is correlated with lake size: limnetic ecomorphology and diet are typical in larger lakes where midwater zooplankton form a greater share of available resources (Ishikawa et al. 2019). These ecotype differences affect stickleback nutrition (Ishikawa et al. 2021), gut microbiota (Bolnick et al. 2014), and parasite infections (Bolnick et al. 2020; Bolnick et al. 2026), which all might modify the metabolome.

Here, we present evidence confirming among-population variation and ecotype divergence in the stickleback liver metabolome. We focus on the liver metabolome because it is involved in over 500 physiological processes related to diet, metabolism, detoxification, and immunity (Yasaka et al. 2026). To partition the role of genotype and environment in these population differences, we take advantage of a large-scale translocation experiment that redistributed stickleback ecotypes across different environments (Hendry et al. 2024). Translocation experiments have been widely used in evolutionary ecology for nearly a century (Clausen et al. 1940) to test for local adaptation and plastic versus evolved population differentiation. To the extent that phenotypic variation is heritable, traits may continue to reflect translocated population’s ancestry, regardless of their current habitat. If variation is plastic, a given ancestry will exhibit different traits in distinct habitats. We show that several generations after the translocation experiment, stickleback metabolomes reflect both their ancestry and their present habitat.

## METHODS

### Experimental design

In 2018, the Alaska Department of Fish and Game used Rotenone in nine lakes on the Kenai Peninsula, Alaska, to eliminate invasive fish species. In June 2019, over 9,000 native stickleback from eight “source lakes” in the Cook Inlet region were trapped and translocated into the fishless lakes to re-establish stickleback populations (see Hendry et al. 2024 for sample sizes, maps, and other details). The source populations are genetically divergent (Bolnick et al. 2026), reflecting approximately 15,000 years of allopatry following Pleistocene deglaciation (Reger and Pinney 1996). The populations vary morphologically from relatively benthic to relatively limnetic (four lakes each; Haines et al. 2023; Hendry et al. 2024). A roughly equal number of fish from the four limnetic source lakes were mixed and introduced into four fishless lakes (two small benthic habitats, two large limnetic habitats, Figure 1). Likewise, a pool of fish from the four benthic sources were mixed and introduced into two small and two large lakes. A ninth lake received both benthic and limnetic fish. One introduction failed (G Lake, benthic ecotypes introduced into a limnetic lake), and was reintroduced in 2022 using benthic and limnetic founders. This transplant design creates three sets of genotype groups: the benthic colonist pool - 0% limnetic ancestry, a mixed pool (50% limnetic ancestry), and the limnetic colonist pool (100% limnetic ancestry), distributed across both large and small lakes, thereby allowing us to statistically partition effects of ancestry and environment. This study compares the metabolomes of the source lakes, and then the results of the translocation experiment.

**Figure 1.**
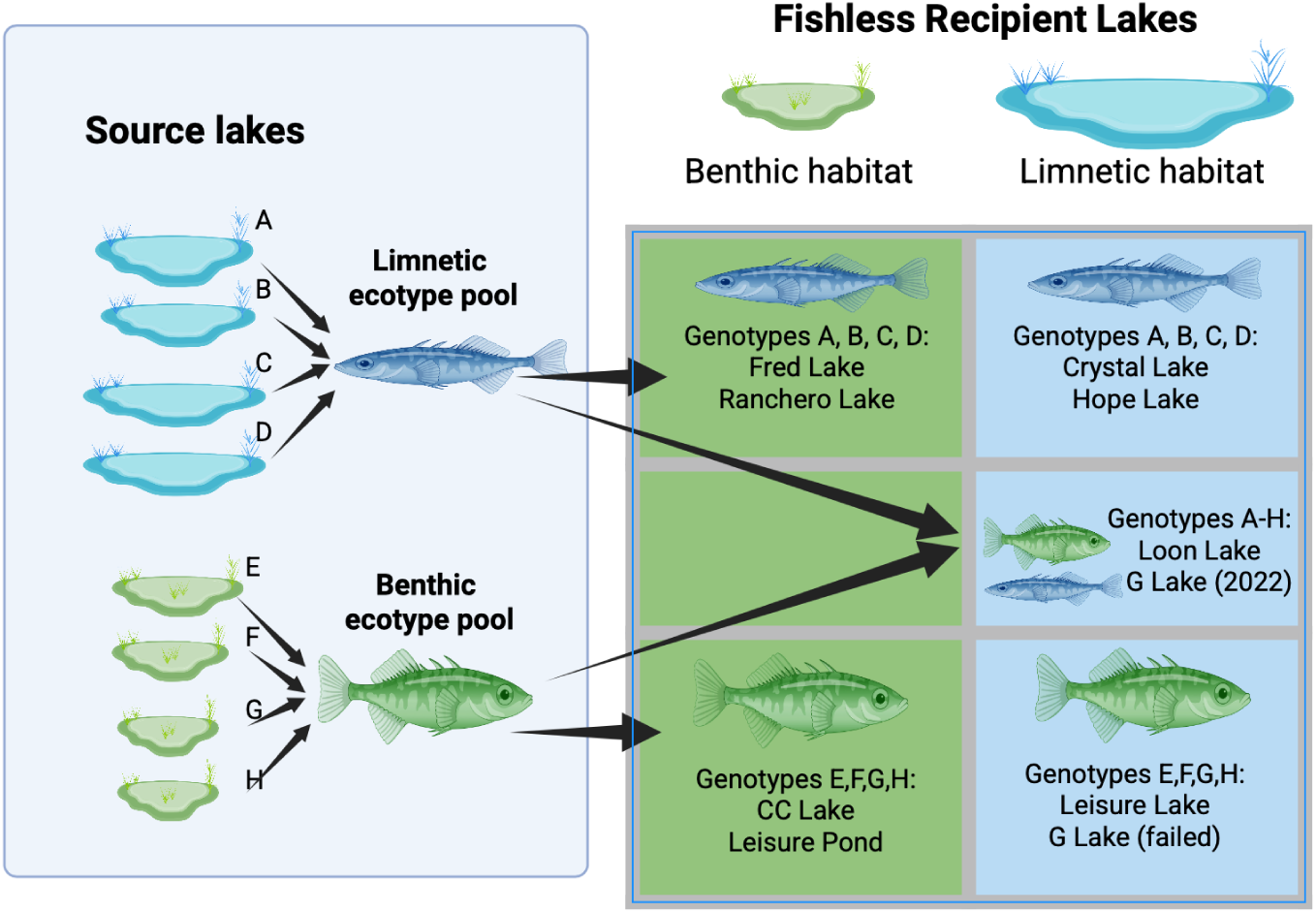
Summary of the experimental design. Source lakes indicate the original resident populations which were selected to provide fish for translocation. Recipient lakes indicate the restoration lakes. The colors of the fish icons refer to the ecotype of the populations based on morphology (Hendry et al. 2024). The table showing the translocation factorial design uses color coding of the table cells to denote lake habitat (relatively small benthic or large limnetic environments). The 2019 introduction to G Lake failed by 2020, and a benthic-limnetic mixture of fish were reintroduced in 2022. Figure generated with BioRender.

### Sample Collection

In late May and early June 2023, we used minnow traps set overnight to sample stickleback from seven source lakes and nine recipient lakes. One source lake population (Long Lake) had been extirpated by invasive Northern Pike and could not be resampled. Stickleback were euthanized in MS-222 within hours of capture. Fresh liver samples were collected into 95% ethanol (Zuffa et al. 2025) for 30 individuals per population (480 individuals total). From these same fish, we recorded anatomical sex by dissection to inspect gonads, and standard length, as well as other traits that will be the subject of later papers (infection, diet, microbiome, transcriptome, genotypes). Permits were provided by the Alaska Department of Fish and Game (SF2023-030d), Alaska State Parks (#2301803), and National Wildlife Refuge (2023-Res-DBolnick-11109). Vertebrate research was approved by the University of Connecticut IACUC (A22-006).

### Extraction

Tissue samples were prepared for LC-MS/MS as described in Caraballo et al. (2024). Briefly, the entire portion of preserved liver tissues in 95% ethanol were dried in a centrifugal vacuum concentrator, Centrivap (Labconco, Kansas City, MO, USA). Rather than standardize the liver to a fixed mass of tissue, we use fish body mass as a covariate in many of the subsequent data analyses. Dried tissue was transferred to a clean centrifugation tube, a 5 mm stainless steel bead and 1 mL of extraction solvent (50% methanol:water, LC-MS grade containing 1uM of sulfadimethoxine as internal standard) per tube were added. Tubes were homogenized at 30 Hz for 8 min in a Qiagen TissueLyzer II. Samples were kept at 4 °C for 1 h for protein precipitation and a centrifugation step was performed for 5 min at 14,000 RPM (Centrifuge 5418 Eppendorf USA). A total of 400 uL of supernatant were transferred to a new 96-well plate and dried out in a centrifugal vacuum concentrator. Prior to LC-MS/MS analysis, dry samples were reconstituted in 200uL of 50% methanol/water (LC-MS grade). A pooled sample (PoolQC) was generated by collecting 10 μL from each biological sample, combining these into a single mixed sample, and aliquoting 200 μL into one well on each plate.

### LC-MS/MS acquisition

Untargeted LC-MS/MS acquisition was performed on a Vanquish Flex UHPLC system coupled to an Orbitrap Exploris 240 (Thermo Fisher Scientific, Bremen, Germany). Chromatographic separation was performed on a Kinetex 2.6 um 100 A pore size Polar C18 reversed phase UHPLC column 100x2.1 mm (Phenomenex, Torrance, CA) with a constant flow rate of 0.5 mL/min. The following solvents were used during the LC-MS/MS acquisition: water with 0.1% formic acid (v/v), Optima™ LC/MS Grade, Thermo Scientific™ (solvent A) and acetonitrile with 0.1% formic Acid (v/v), Optima™ LC/MS Grade, Thermo Scientific™ (solvent B). After injection of 2μL of sample into the LC system, the elution was performed isocratically with 5% B from 0-1.0 mins, then with a linear gradient from 5-99% B (0.5-6.0 min), 99% B (6.0-7.5 min), 99-5% B (7.5-8.0 min), 5% B (8.0-8.5 min), 5%-99% B (8.5-8.75 min), 99% B (8.75-9.25), 99%-5% B (9.25-9.5 min) and 5% B (9.5-10.0 min).

Data dependent acquisition (DDA) mode was used for acquisition of tandem MS (MS/MS) data with a default charge state of 1. Full MS was acquired using 1 microscan at a resolution (R) of 120 000 at m/z 200, normalized automatic gain control (ACG) target 100%, maximum injection time (IT) of 100 ms, scan range m/z 100-1,100 and data acquired in profile mode. DDA of MS/MS was acquired using 1 microscan at a resolution (R) of 15,000 at m/z 200, normalized automatic gain control (ACG) target 100%, top 10 ions selected for MS/MS with isolation window of m/z 1.0 with scan range m/z 100-1,100, fixed first mass of m/z 50 and HCD collision energies (%) of 20, 30, 40 and 50. Analytical blanks, pool samples (PoolQC) and mixture of sulfamethazine, sulfamethizole, sulfachloropyridazine, sulfadimethoxine, amitriptyline, and coumarin-314 (QCmix, 10 μM) were injected after every 10 samples as quality control for monitoring instrument (LC-MS) performance. Injection order was randomized to minimize variation during data acquisition.

### LC-MS/MS data processing, quality control, and normalization

Raw LC-MS/MS data files were processed in MZmine version 4.7.29 (Schmid et al. 2023). Preprocessing steps were performed to generate a feature table and peak list (Heuckeroth et al. 2024) for use in the GNPS2 platform (Wang et al. 2016), in addition to an mgf file for SIRIUS annotations (Dührkop et al. 2019).

Blanks, internal standard and QCmix samples were excluded from downstream analysis in R version 4.4.2. Feature retention was determined based on whether the pool to blank ratio and the pool to QCmix had a ratio greater than ten. Features with variances near zero were removed using the caret package version 7.0.1 in R and the associated nearZeroVar function (Kuhn, 2008). We also excluded several known contaminants (the anesthetic MS-222 used to euthanize the fish, and DEET used by field personnel as insect repellent). This retained a matrix with 5,939 features, which we normalized using rCLR to obtain relative metabolite abundance.

Normalization was computed using the log of each feature intensity divided by the geometric mean of non-zero features within each sample; this is a typical normalization for LC-MS data (Martino et al. 2019).

### Characterization of the stickleback metabolome

To characterize the chemical composition of the stickleback liver metabolome, a feature-based molecular network (FBMN) was created using the online workflow (https://ccms-ucsd.github.io/GNPSDocumentation/ ) on the GNPS2 website (https://gnps2.org/ ). The data was filtered by removing all MS/MS fragment ions within +/- 17 Da of the precursor m/z. MS/MS spectra were window filtered by choosing only the top 4 fragment ions in the +/-50Da window throughout the spectrum. The precursor ion mass tolerance was set to 0.02 Da and a MS/MS fragment ion tolerance of 0.02 Da. A network was then created where edges were filtered to have a cosine score above 0.7 and more than 4 matched peaks. Further, edges between two nodes were kept in the network if and only if each of the nodes appeared in each other’s respective top 10 most similar nodes. Finally, the maximum size of a molecular family was set to 100, and the lowest scoring edges were removed from molecular families until the molecular family size was below this threshold. The spectra in the network were then searched against GNPS2’ spectral libraries. The library spectra were filtered in the same manner as the input data. All matches kept between network spectra and library spectra were required to have a cosine score above 0.7 and at least 4 matched peaks. The molecular networking analysis with default and propagated libraries of bile acids (Wang et al. 2016) can be accessed through the link: https://gnps2.org/status?task=a78a3532e6f14df38f61183b0835ed53.

From the GNPS spectral library, 1,469 chemical annotations were generated that had a minimum cosine score of 0.7 and a minimum of 4 matched peaks. To assign annotations to chemical compounds that were not identified by GNPS, SIRIUS version 6.3.3 was run to generate formulas with CSI:FingerID for structure associations against other metabolite databases (Dührkop et al. 2019). Chemical compound annotations were broken into three tiers based on the Metabolomics Standard Initiative (MSI), which is a set of reporting standards that consider the methodology used to generate annotations (Sumner et al. 2007). MSI Level 1 classification is given to chemical standards, MSI Level 2 classification is given to features that had a GNPS spectral library match (cosine score > or equal to 0.7 and > or equal to 4 matched peaks), and MSI Level 3 classification is given to features that had structural predictions from SIRIUS CSI: FingerID or chemical class predictions from CANOPUS NPCLassifier (Sumner et al. 2007; Schymanski et al. 2014). CANOPUS NPCLassifier was run with SIRIUS to generate chemical classes without requiring databases or a structural match from CSI: FingerID (Dührkop et al. 2021; Kim et al. 2021). All structural annotations from this study are putative, as no chemical standards were used.

The generated, annotated molecular network from GNPS was visualized in Cytoscape version 3.10.4. (Shannon et al. 2003). All connected nodes are shown in the full network and were assigned a color based on the NPClassifier chemical class assignments made through CANOPUS (Kim et al. 2021).

Spectral similarity searches against all public mass spectrometry datasets were conducted using https://fasst.gnps2.org/fastsearch/ and the metabolomicspan_index_nightly library (Pullman et al. 2021; Bittremieux et al. 2020; Wang et al. 2018). Results were grouped by dataset and inspected for matches in fish, liver, or stickleback-specific studies by cross-referencing dataset accession numbers on <u>massive.UCSD.edu</u>.

Overrepresentation analysis (ORA) of CANOPUS NPClassifier chemical classes was conducted using a right-tailed fisher’s exact test with Benhamini-Hochberg FDR correction in R version 4.42 (Wieder et al. 2021). A background was generated using only detected and retained features with NPClassifier class annotations (n=3,443 features). Enrichment was tested at the class-level using a log Fold-Change (lFC) threshold of 0.3 as the foreground filter, with sensitivity assessed at lFC thresholds of 0.2 and 0.5. ORA was run on exploratory individual-level Limma features as a hypothesis-generating class-level tier of evidence.

### Analysis: Does the stickleback metabolome vary among source lake populations?

We conducted Principal Component Analysis (PCA) of the rCLR-normalized feature matrix to summarize metabolomic variation among individuals, focusing initially on source lake samples (N=210 fish). To test for differences among the source lake populations, we used multivariate analysis of variance (MANOVA) on the first 100 principal components (PCs), with covariates sex and standard length, and all interactions. Results were robust to the choice of numbers of PC axes. For a supervised ordination, we used discriminant function analysis of principal components (DAPC; Jombart et al. 2010), trained to separate fish by population. All DAPC analyses retain a number of PCs based on optimal α score. Classification accuracy is reported as resubstitution accuracy. We used Type III sums of squares ANOVAs to separately test each of the top 25 PC axes for effects of lake, sex, and standard length, and their interactions.

To identify metabolites that are characteristic of individual populations, we performed a one-vs-all lake contrast analysis using Limma package version 3.62.2 (Ritchie et al. 2015) with empirical Bayes variance shrinkage, comparing each lake mean against the mean of all other lakes. All P-values from these analyses were adjusted using a Benjamin-Hochberg False Discovery Rate (FDR) threshold of<0.05.

### Analysis

#### Is among-population metabolomic variation driven in part by geography or ecotype?

To test for geographic contributions to the metabolomic variation, we classified each lake by its region (Matsu Valley or Kenai Peninsula) and used a MANOVA to test for between-lake differences as a function of region. To avoid pseudoreplication we use lakes as the level of replication (N=7), because replicate fish from the same lake provide non-independent information on how lake-level characteristics (e.g., geography, population mean ecomorphology) affect the metabolome. Therefore the MANOVA was applied to population centroids (PCA means) for the leading 5 PC axes.

To test the effects of population ecotype on the metabolome, we also used a MANOVA applied to the centroids of PC1-5. For supervised learning, we used DAPC to assess the separability of source lakes based on ecotype identity. We used a linear model to test for an ecotype effect on population mean discriminant axis 1 (DA1) values (N=7 lakes).

To explore relationships between individual metabolites and population ecotype, we assessed differential feature abundance using a nested linear model strategy in which the population is a blocking variable nested within ecotype, using individual sex and standard length included as covariates. The consensus intra-lake correlation was estimated using the duplicateCorrelation function and incorporated into the linear model via Generalized Least Squares (GLS) accounting for non-independence of individuals within the same lake. To independently cross-validate the differential abundance results, we performed a partial least squares discriminant analysis (PLS-DA) using the mix0mics package . PLS-DA is primarily used to extract variable importance in projection (VIP) scores for each feature. Features with VIP > 1 on the first component were used as an independent foreground set to confirm which features and chemical classes strongly differentiate ecotypes across two frameworks. Chemical class enrichment was assessed separately for features elevated in benthic source fish (logFC>0.3) and features elevated in limnetic source fish (logFC<-0.3) using CANOPUS ORA.

### Analysis

#### Does metabolome variation in recipient populations reflect ancestry or habitat?

Data processing, statistical analyses, and graphics were performed in R version 4.42 (R Core Team, 2024). Using fish sampled from the nine experimental recipient lakes (N=270), we estimated the metabolomic effects of ancestral ecotype, and present-day habitat. We applied PCA to the rCLR-normalized feature matrix for recipient lake fish only. We used a MANOVA to test for effects of ancestry and lake habitat on the population centroids for PC axes 1-5 (lakes represent the level of replication). We coded ancestry in two ways: first, as a factor with three levels (benthic, limnetic, or mixed); second, using a quantitative scale (0%, 50%, 100% limnetic). The latter approach has greater power to detect additive genetic effects, if the mixed-founder lakes are intermediate; although subsequent evolution modified the exact ancestry proportions in the mixed lakes (Eckert et al. 2026). We used Type III ANOVAs to test whether each leading PC axis depended on ancestry and lake habitat, using lake means as the dependent variable. We used DAPC for supervised classification, training the model on combinations of ancestry and habitat. To test classification success, we calculated lake means for discriminant axes DA1-DA3, and tested whether these depend on lake ancestry, habitat, and their interaction.

The experimental design generated populations that were either pre-adapted (e.g., limnetic ecotype pool added to limnetic lakes) or maladapted (e.g., benthic fish introduced to limnetic lakes) to their new habitats. To test for metabolomic effects of this ecotype-habitat mismatch, we reran DAPC trained to separate recipient lake fish by the combination of their ecotype, and whether the ecotype matched the habitat. We tested for effects of ecotype, mismatch, and their interaction, in both multivariate space (MANOVA applied to lake centroids for DA1-3) and each discriminant axis separately (Type III ANOVAs).

To identify the particular metabolome features underlying ecotype differences in recipient lakes, we used differential-abundance contrasts in nested linear models, as described above for source lakes. We ran two contrasts, first modelling the effect of the benthic compared to limnetic ancestry, then modeling the effect of habitat. Chemical class enrichment through ORA were applied to the blocked differentially abundance results using a logFC threshold of 0.3 and a CANOPUS NPClassifier background of 3,443 features without filtering out for p-value significance, as described for source lakes. These ORA results are treated as hypothesis-generating class-level evidence to observe chemical trends. Following generation of chemical trend patterns using ORA, the annotated features with significant changes in recipient fish, specifically features reduced in recipient fish in the destination contrast, were filtered for the putative classes and chemicals given by ORA. Molecular family membership was verified by network visualization using Cytoscape.

## RESULTS

### Characterization of the stickleback metabolome

Untargeted LC-MS/MS profiling of 480 stickleback from 16 lakes yielded 23,371 detected features with 5,939 features retained after quality control filtering (Supplementary Figure 1). Of these, 1,469 received MSI Level 2 annotations through GNPS spectral library matching, 358 received MSI Level 3 structural predictions through SIRIUS CSI:FingerID, and 2,667 received chemical class predictions through CANOPUS NPClassifier (Supplementary Figure S1). The annotated chemicals composing the stickleback liver metabolome were dominated by fatty acids (n=1,728 chemically distinct compounds, 50.2% of CANOPUS-annotated features) and amino acids and peptides (n=1,139, 33.1%). The remainder was comprised of polyketides (n=284, 8.2%), carbohydrates and terpenoids (n=94, 2.7%), alkaloids (n=57, 1.7%), and shikimates and phenylpropanoids (n=47, 1.4%) (Figure 2). Chromatographic separation confirmed expected chemical space placements, with polar metabolites including amino acids eluting at retention times of 2 to 4 minutes and nonpolar lipids eluting at 5 to 7 minutes on the reverse-phase C18 column (Supplementary Figure S2). Fatty acids had the largest proportion of MSI Level 2 annotations of any chemical class reflecting good spectral library coverage for lipid classes, while polyketides, terpenoids, alkaloids, and shikimates were predominantly MSI Level 3 (Supplementary Figure S3).

**Figure 2.**
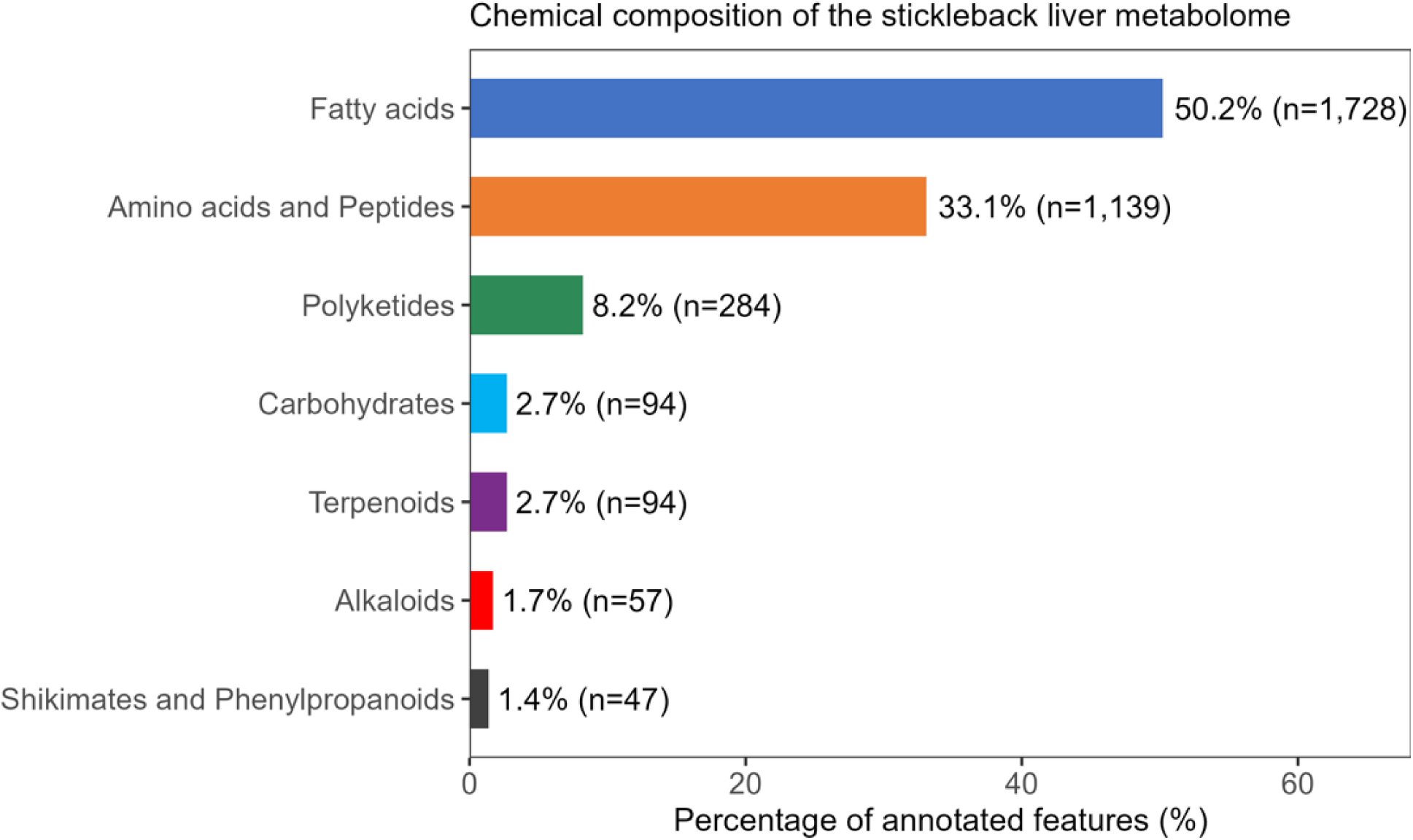
Bar chart showing chemical class composition of the threespine stickleback liver metabolome. Denoted by color, the classes are fatty acids (blue), amino acids and peptides (orange), polyketides (green), carbohydrates (blue), terpenoids (purple), alkaloids (red), and shikimates and phenylpropanoids (black). Bar size reflects the abundance of features per chemical class predicted by CANOPUS NPClassifier (n=3,443 features with class-level predictions, MS-222 and DEET excluded). All annotations here are putative (MSI Level 3).

### The stickleback liver metabolome differs among source populations

The source lake populations of stickleback differ significantly in their liver metabolomes, despite considerable overlap along any one axis (Figure 3). PCA of the normalized feature matrix suggests that biological variation in the metabolome is high-dimensional, with variance distributed across many PC axes with modest eigenvalues (PC1=12.4%, PC2=4.6%, PC3=3.1%, PC4=2.8%, PC5=2.3%). The top 100 PCs cumulatively explain 74.8% of variance. MANOVA analysis of these individual PCA scores reveals that sex is the largest source of variation among source lake fish (Supplementary Figure S4; Pillai’s trace **λ**=0.88, df=1, P<0.0001, partial η^2^=22.0% variance explained), with lake close behind (Pillai’s trace **λ**=4.8, df=6, P<0.0001, η^2^=20.0%), followed by length (η^2^=16.1%, P<0.0001), lake*sex interaction (η^2^=13.5%, P<0.0001), and lake*length interaction (η^2^=9.7%, P<0.0001). Similarly, univariate ANOVAs of leading PC axes consistently find evidence for among-lake variation. For PC1, source lake is the strongest effect (F_6,182_=9.7, P<0.0001), with a modest contribution of fish length (F_1,182_=6.54, P<0.011) and a lake*sex interaction (F_6,182_=2.5, P<0.023). Lakes also differed significantly along PC2 (F_6,182_=5.0, P<0.0001), PC3 (F_6,182_=21.5, P<0.0001), and almost all of the top 25 PC axes, most of which also exhibit lake*sex or lake*length interactions. Diagnostics (e.g., qq-plots) suggest reasonable fit to linear model assumptions.

**Figure 3.**
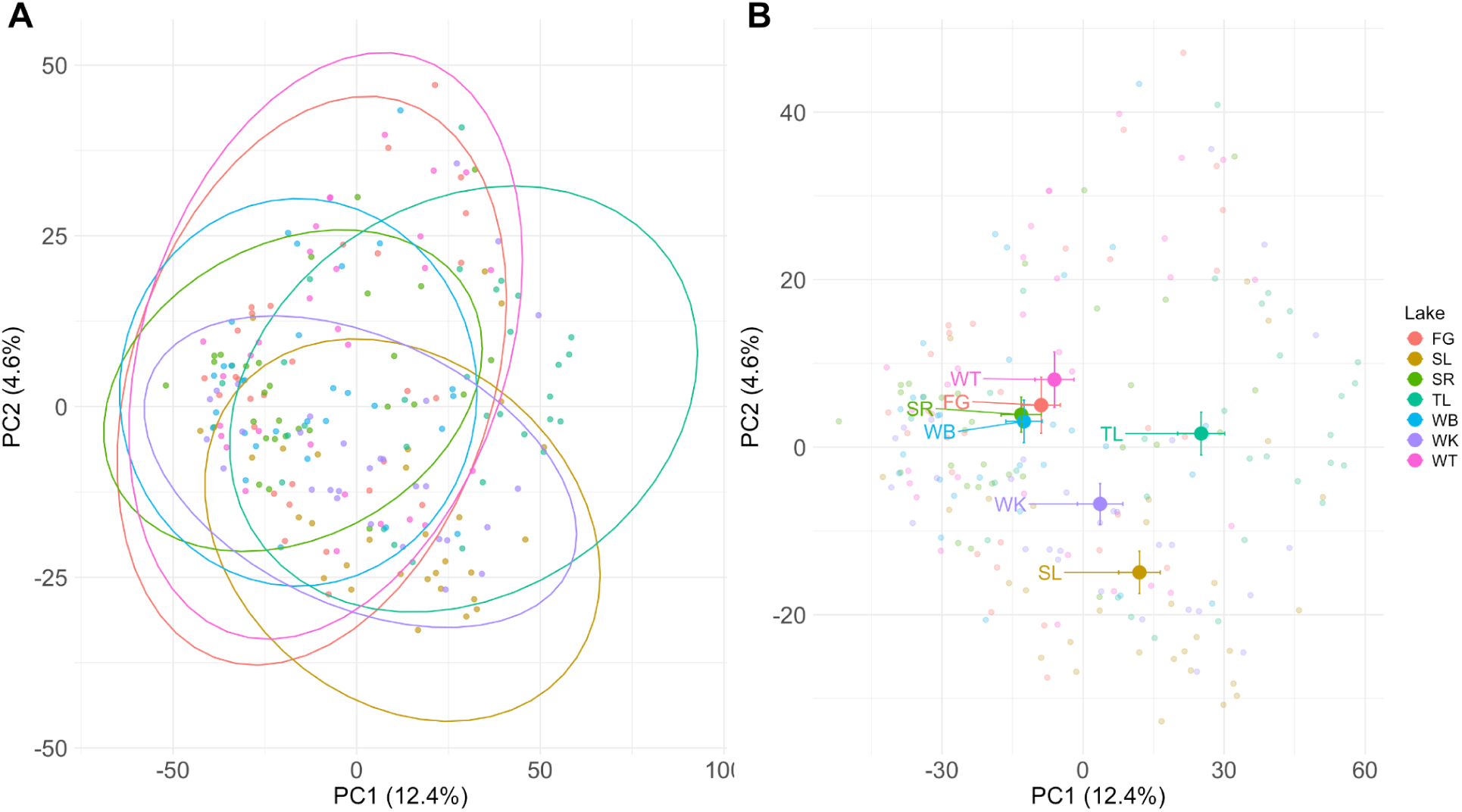
PCA results on the rclr-transformed dataframe for the source lakes. PC1 and PC2 explain less than 15% of the global variance, and source lakes overlap considerably in PC composition (at least in this 2-D projection). A) PC1 vs PC2 with lake 95% ellipses for each population, where points are individual fish positions on the PCA. B) PC1 vs PC2 with lake population means (centroids) with 1 standard error bars to indicate confidence in between-population differences in average metabolomes. Lake abbreviations: FG = Finger Lake, SL = Spirit Lake, SR = South Rolly Lake, TL = Tern Lake, WB = Walby Lake, WK = Wik Lake, WT = Watson Lake.

Despite this strong support for differences in population means, there is clearly extensive overlap between populations attributable to sex, size, and likely other sources of variation not accounted for here. As a result, discriminant analysis classification success is modest. If we use only PC1 and PC2 in DAPC, 34% of fish can be correctly classified to their lake of origin (low, but better than random ***χ***^2^=32.2, P<0.0001). However, using more PC axes increases discriminant analysis classification success (72% accuracy with 10 PCA axes, or 83% with 20 axes).

Of the 5,939 retained features, 4,864 (81.9%) were detected in all seven sampled source lakes, and no features were unique to a single source lake, suggesting that metabolomic differences among populations are quantitative (abundance) rather than qualitative (presence/absence). By testing each metabolite feature separately for variation among source lakes, 2,407 features were significantly elevated in at least one source lake relative to others (FDR<0.05), confirming that lake of origin is an important driver of liver metabolomic variation in stickleback. These lake effects can arise from genetic differences between lakes, or environmental differences (abiotic water conditions, climate, microbiota, diet, parasitism, etc). The most variable annotated features across source lakes included fatty acids and phospholipids, which were most abundant in Tern and Spirit Lakes.

### Geography and ecotype contribute to variation among source lakes

Metabolomic variation among source lakes partly reflects their geography. For instance, in Figure 3B, the tight cluster of centroids are mostly Mat-Su valley lakes, whereas the three in the lower right are Kenai Peninsula lakes. A MANOVA on the PC1-25 lake centroids confirms a modest effect of geography (P=0.05). Using supervised learning (DAPC) to separate individuals by region, the discriminant axis readily separates regions (F_1,5_=19.71, P=0.007; Supplementary Figure S5).

The stickleback ecotypes (benthic vs limnetic populations) also differ in metabolomic composition. Unsupervised ordination (PCA) showed no significant ecotype effect in a MANOVA applied to population centroids (N=7 lakes). However, in univariate ANOVAs, PC5 provided strong support for an ecotype effect (F_1,4_ =44.75, P=0.0026), even after Bonferroni correction for testing 10 PC axes. Supervised learning methods (DAPC) provide better ecotype resolution (Figure 4): an ANOVA of DA1 lake means reveals significant effect of ecotype (F_1,5_=63.73, P=0.0005), with high classification success for individual fish. Notably, population ecotype (based on morphology; Hendry et al 2024) is more effective at supervised learning, compared to lake surface area (ANOVA of DAPC axis 1, F_1,5_=16.7, P=0.01).

**Figure 4.**
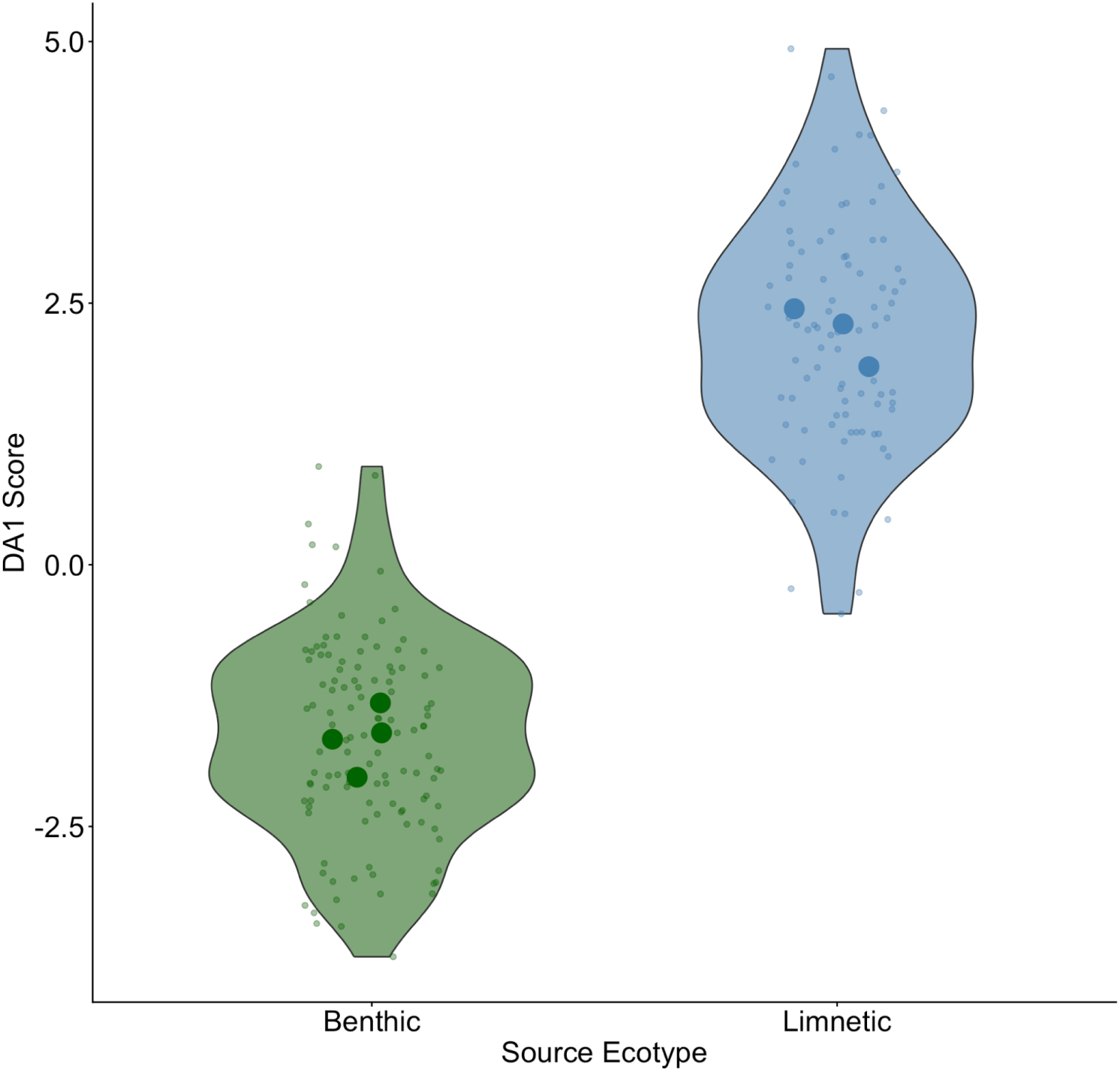
Separation of source lake limnetic and benthic ecotype populations by discriminant analysis of principal components (DAPC) using PC axes 1-25. Individual fish are represented as small points, population centroids as larger circles color coded by ecotype.

Differential abundance analysis and VIP scores from PLS-DA both identified acylcarnitines as the primary chemical class distinguishing limnetic from benthic ecotypes in source lakes. Exploratory individual-level differential abundance analysis identified 61 features elevated in one ecotype relative to the other in source lakes, with 12 elevated in benthic and 49 elevated in limnetic fish. Although none of these features are significant after false-discovery rate correction, there are too many uncorrected-significant effects to be error alone: a proportion test confirmed that 9.1% of features showed nominal ecotype differences, significantly exceeding the 5% expected under the null (p<0.0001). This suggests that there is a high-dimensional metabolic signal of ecotype, even though no single feature captures this robustly at the among-lake level.

Two different analytical approaches identified acylcarnitines as the primary metabolite class distinguishing limnetic from benthic fish: multivariate classification via PLS-DA identified acylcarnitines as significant VIP features; also, differential abundance analysis identified 2 acylcarnitine features with FDR<0.05 (features 21087 and 12275), with several other features trending in the same direction. Among the most consistently elevated annotated features in limnetic source fish in the differential abundance analyses were hexanoylcarnitine (logFC=-0.772), pentanoylcarnitine (logFC=-0.888), butyrylcarnitine (logFC=-0.540, not FDR significant), and acetylcarnitine (logFC=-0.326, not FDR significant). These features were annotated at MSI Level 2 through GNPS library matching (Figure 5A).

**Figure 5.**
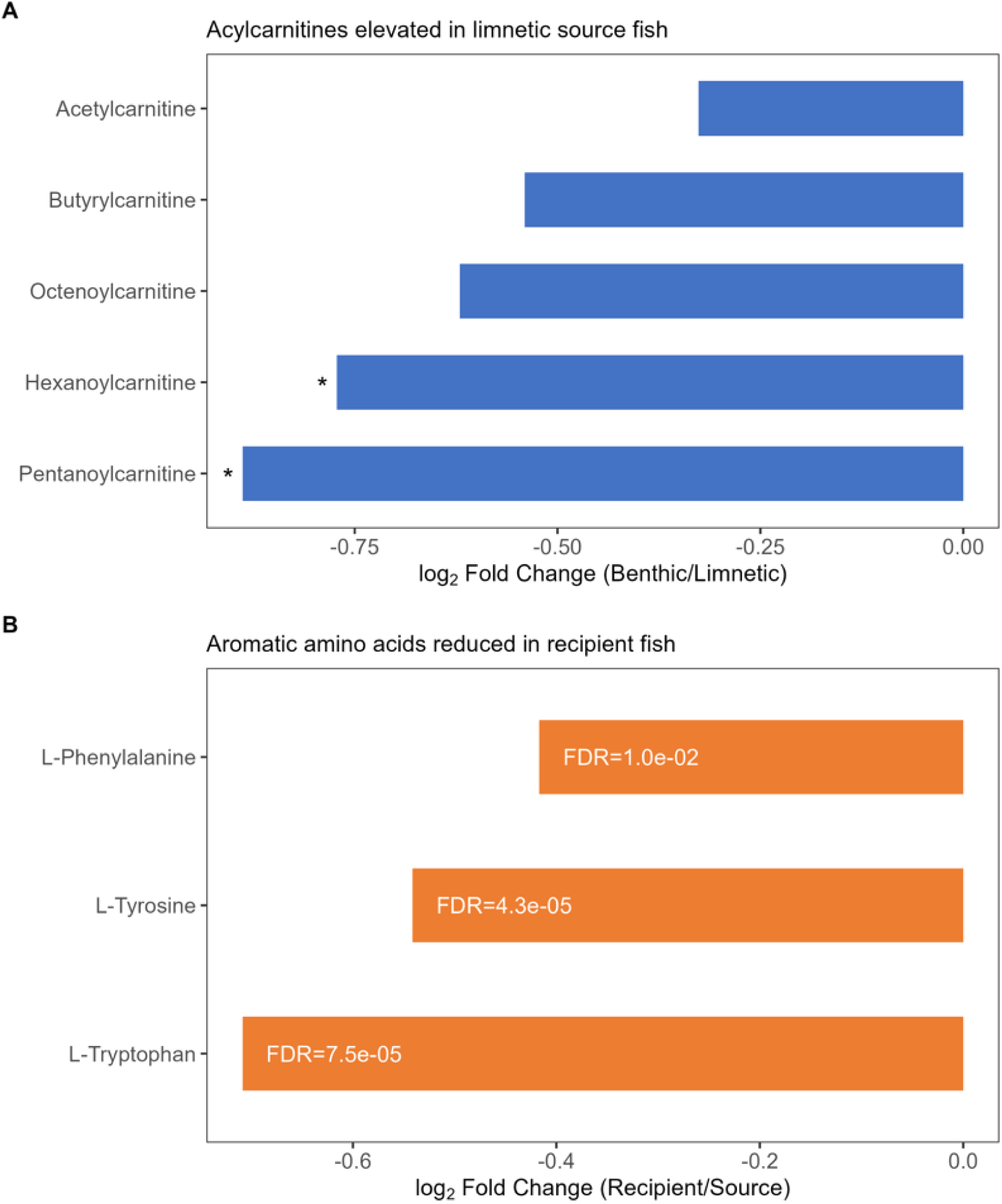
Annotated metabolite features elevated in specific biological contrasts. A) Acylcarnitine features elevated in limnetic source fish relative to benthic source fish (source ecotype contrast; lFC<0=higher in limnetic; MSI Level 2 GNPS spectral match). B) Aromatic amino acids elevated in source fish relative to recipient fish across both ecotypes (destination contrast; lFC<0=higher in source; MSI Level 2 GNPS spectral match; all three are also elevated in limnetic source fish, FDR<0.05).

Multiple taurocholic acid spectral cluster features had VIP > 1 in the ecotype PLS-DA model (comp1 range 1.01-1.63), and molecular networking confirmed these features as part of a cholane steroid spectral cluster (Figure 6A). Likewise, acylcarnitines with high VIP scores formed a spectral cluster (Figure 6B). Additionally, fastMASST spectral search detected taurocholic acid feature 32963 in two other stickleback liver datasets (Supplementary Figure S6).

**Figure 6.**
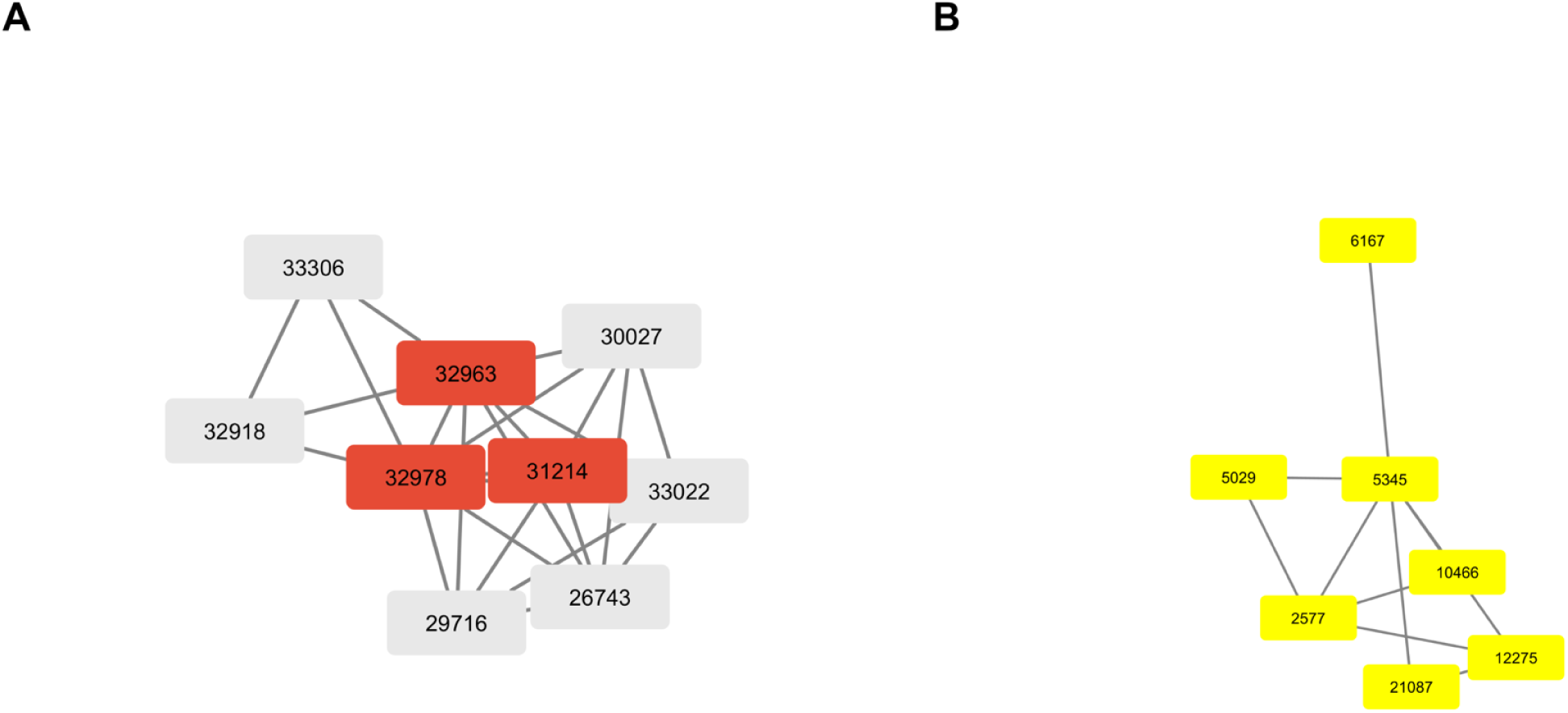
Molecular network visualization of key annotated metabolite clusters using Cytoscape (Wang et al. 2016). A) First-neighbor spectral network of taurocholic acid feature 32963. Red nodes indicate CANOPUS cholane steroid class annotation. Edges represent cosine similarity > or equal to 0.7 and > or equal to 4 matched peaks. Feature 32963 is within a cholane steroid molecular family. B) Yellow nodes indicate acylcarnitine features significantly elevated in limnetic source fish (FDR<0.05). Edges indicate high cosine similarity between acylcarnitine nodes.

### Ancestry and habitat jointly shape metabolomic variation in recipient populations

The translocation experiment changed the nature of benthic-limnetic ecotype differences (Figure 7A). Analyzing fish from all 16 lakes in a DAPC trained on lake category (source vs. recipient) and ecotype (benthic, limnetic, or mixed), reveals clear separation between the source and recipient lakes (DA1 lake category F_1,12_=183.4, P<0.0001), which may reflect effects of new habitat, genetic admixture, stress, or ecological opportunity in underpopulated lakes. Benthic versus limnetic source lakes are separated by DA2 (ecotype F_1,12_=66.7, P<0.0001), but the recipient lakes cluster tightly in this axis, resulting in a lake category*ecotype interaction (F_1,12_=31.8, P=0.0001). Thus, the metabolomic features that separate benthic versus limnetic source lakes are uninformative regarding the features that separate recipient lake ancestries (discriminant axis cosine similarity r = 0.13, angle=82.36°).

**Figure 7.**
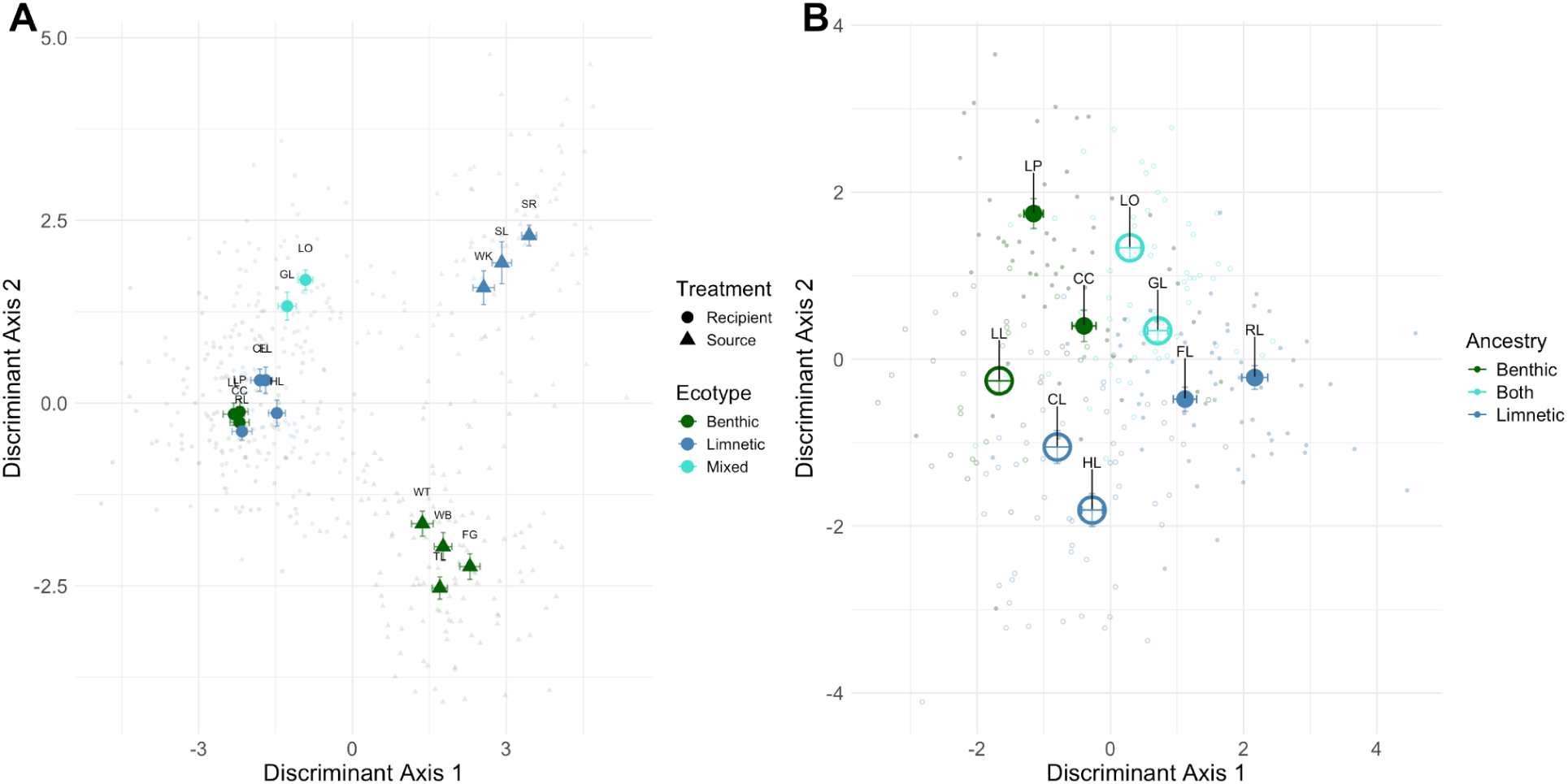
A) Discriminant analysis of principal components (DAPC) applied to all sampled fish (source and recipient), trained to separate both lake category and population ecotype. Individual fish data points are given as small dots, population centroids (the relevant level of replication) are large points. Color coding distinguishes the ecotype of fish used to found each population. Point types distinguish between source versus recipient lakes. Each centroid is represented with one standard error bars. B) DAPC applied to only recipient lake fish to separate both founder ecotypes (ancestry) and lake habitat (small versus large lakes, represented by smaller filled circles and larger open circles respectively).

Despite this large effect of translocation, we observe effects of both ecotype ancestry and lake habitat on the recipient lake metabolomes. Applying a MANOVA to the centroids of recipient-lake PC axes 1-25 reveals a significant effect of lake habitat (Pillai’s Trace=0.9997, df=1,5, P=0.0314) but not founder ecotype category (Pillai’s=1.31, df=2,5, P=0.6669). When we recode ancestry on a continuous scale (so mixed ancestry is intermediate), the MANOVA is marginally significant for both ancestry (Pillai’s=0.971, df=1,6, P=0.072) and habitat (Pillai’s=0.969, df=1,6, P=0.0765). In univariate analyses, no single unsupervised PC axis yielded significant effects of ancestry or habitat (N=9 lakes as the level of replication).

Supervised learning methods (DAPC) were more effective at separating populations by founder ecotype and lake habitat (Figure 7B), with 77.8% accuracy at assigning fish to the correct founder/lake combination. Lake mean DA1 depended on both founder ecotype (F_2,5=_7.5, P=0.0311) and lake habitat (F_1,5=_13.4, P=0.0145). Lake mean DA2 also differed by founder ecotype (F_2,5=_8.3, P=0.0261) and lake habitat (F_1,5=_6.7, P=0.0491). These inferences were robust to varying numbers of PC axes used for DAPC. If we instead score ancestry on a quantitative scale, the model performance is reduced and only ancestry is marginally significant (P<0.1). Overall, ancestry explained slightly more of the DAPC variation (DA1 44.9%; DA2 58.6%) than did lake habitat (DA1 40.1% and DA2 23.7%). The discrepancy between the unsupervised approach (MANOVA on PC axes) and supervised approach (DAPC) suggests that the latter’s capacity to separate founder ecotypes relies on a combination of the higher-order (small effect) PCA axes.

If metabolomic variation is controlled by additive genetic variation, we would expect that the mixed-ancestry populations would be intermediate between the benthic- and limnetic ecotypes. This is not supported for some DAPC projections (e.g., Fig. 7A), where mixed-ancestry lakes are transgressive. However, when we train a DAPC to separate benthic versus limnetic source lake fish, then apply that discriminant function to recipient lake fish, the axis effectively separates benthic versus limnetic ancestry, and the two mixed-ancestry populations are intermediate (Supplementary Figure S7).

### Metabolomic effect of ecotype-habitat mismatch

The experimental design resulted in some populations whose founder ecotypes were mismatched to their new habitat (e.g., benthic fish added to larger limnetic lakes). Using DAPC trained on ecotype and habitat match, we see effects of this ecotype-habitat mismatch on population means for three of four discriminant axes. Along DA1, mismatched populations diverge in an ancestry-dependent manner (Figure 8A, ancestry*mismatch interaction F_1,3_=11.6, P=0.0420; both main effects P>0.1). DA2 separates benthic and limnetic ecotypes in their matched habitats (Figure 8B; ancestry F_1,3_=15.3, P=0.0296). But, they converge along DA2 when mismatched (ancestry*mismatch interaction F_1,3_=5.7, P=0.0969): limnetic ecotypes become more benthic-like in benthic lakes, and vice versa for benthic ecotypes in limnetic lakes. DA4 shows a directional shift due to mismatch (F_1,3_=22.9, P=0.0173; Figure 8C).

**Figure 8.**
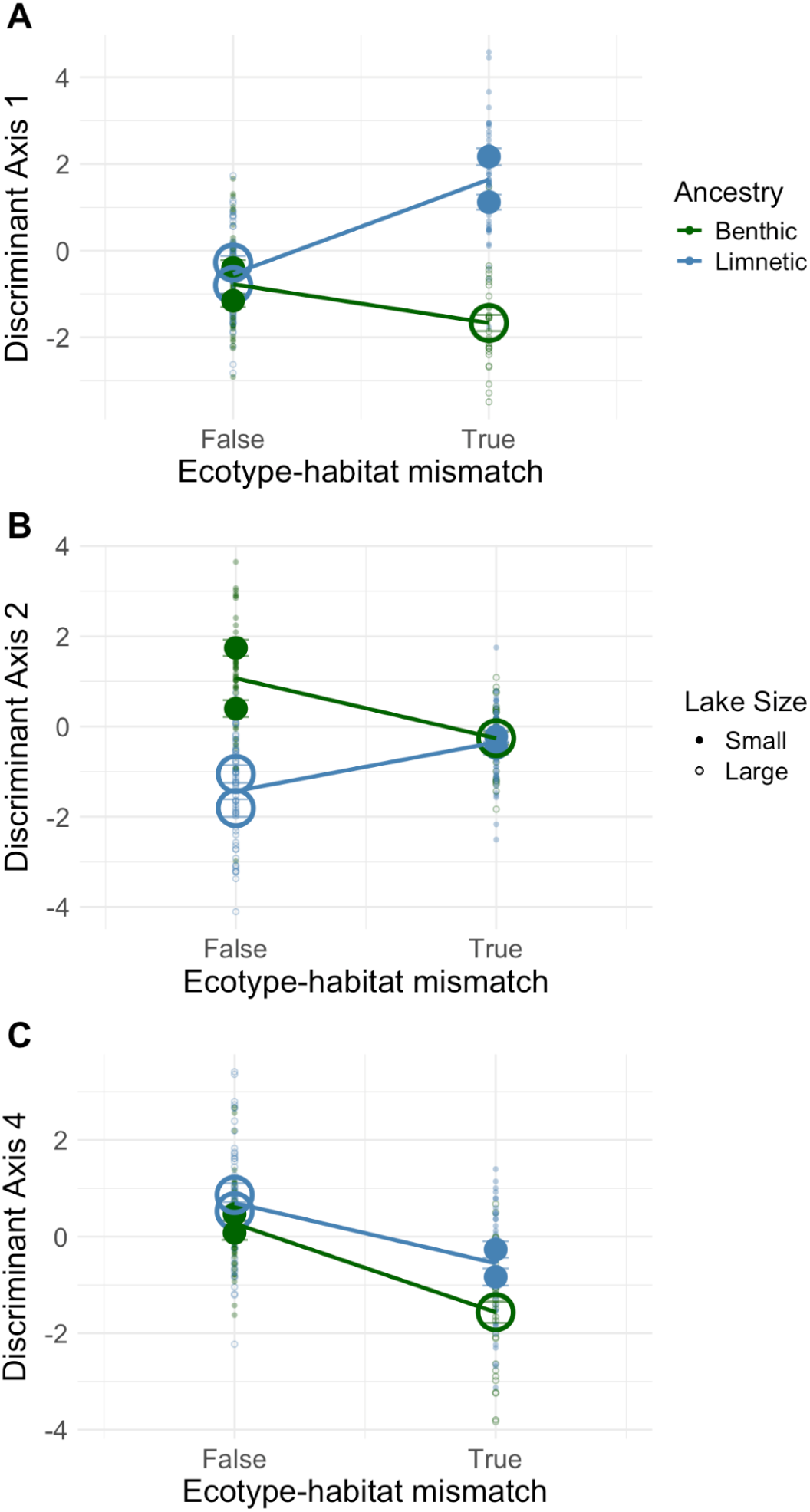
Stickleback populations whose ancestral ecotypes were mismatched to their new lake habitat show distinctive metabolomic profiles. Panels A-C present discriminant axes (1,2, and 4 respectively) obtained from DAPC of recipient lake fish, trained to separate founder ecotype and lake habitat combinations. Individual fish are represented as small points, lake centroids (with error bars) as large points. Points are colored by ancestry (benthic or limnetic ecotype founders added to a lake; mixed-ancestry is omitted here). Small and large lakes (benthic and limnetic habitats) are shown as filled small circles or large open circles. Mismatched populations are either benthic ancestors added to large lakes (green large circles), or limnetic ancestors added to small lakes (blue filled circles).

### Metabolites underlying ancestry and environment effects

To identify candidate annotated compounds underlying the multivariate ancestry signal in recipient lakes, individual-level differential abundance analysis was used. This analysis only identified 23 features nominally elevated in one ancestry group relative to the other (proportion test p=1.00, null result). To disentangle the effects of genetic ancestry from environmental plastic responses within recipient lakes, metabolite abundances were instead modelled using a linear interaction framework, with ecotype and lake size as covariates. Using this interaction model, 31 features were identified that displayed a significant linear ancestry slope, with 12 features associated with benthic ancestry and 19 features associated with limnetic ancestry.

Simultaneously, the habitat of the recipient lake exerted a strong environmental effect independent of ancestry: 77 metabolites were significantly associated with lake size. Of these features, 39 were significantly elevated in large (limnetic) recipient lakes, while 38 were significantly elevated in small (benthic) recipient lakes (Supplementary Figure S8). Twenty features were significantly associated with the ecotype x lake size interaction. Minimal features overlap with those observed as differentially expressed between ecotypes in the source lakes, suggesting they represent plastic metabolic responses to novel habitats rather than ancestry.

Limnetic fish showed greater metabolomic reorganization following translocation than benthic fish. In limnetic fish, 346 feature abundances differed significantly between source and recipient populations (FDR<0.05), with 210 features elevated in recipient fish (potential plastic response), and 136 elevated in source fish (source-associated signal). In benthic fish, only 43 features differed significantly (FDR<0.05), with 19 elevated in recipients and 24 elevated in source fish. This large difference in total affected features indicates that limnetic fish experienced greater metabolomic reorganization following translocation than benthic (Figure 9A). They might either be more metabolomically plastic; or, the recipient lakes may represent a relatively benthic environment requiring greater plastic changes on the part of limnetic source fish.

**Figure 9.**
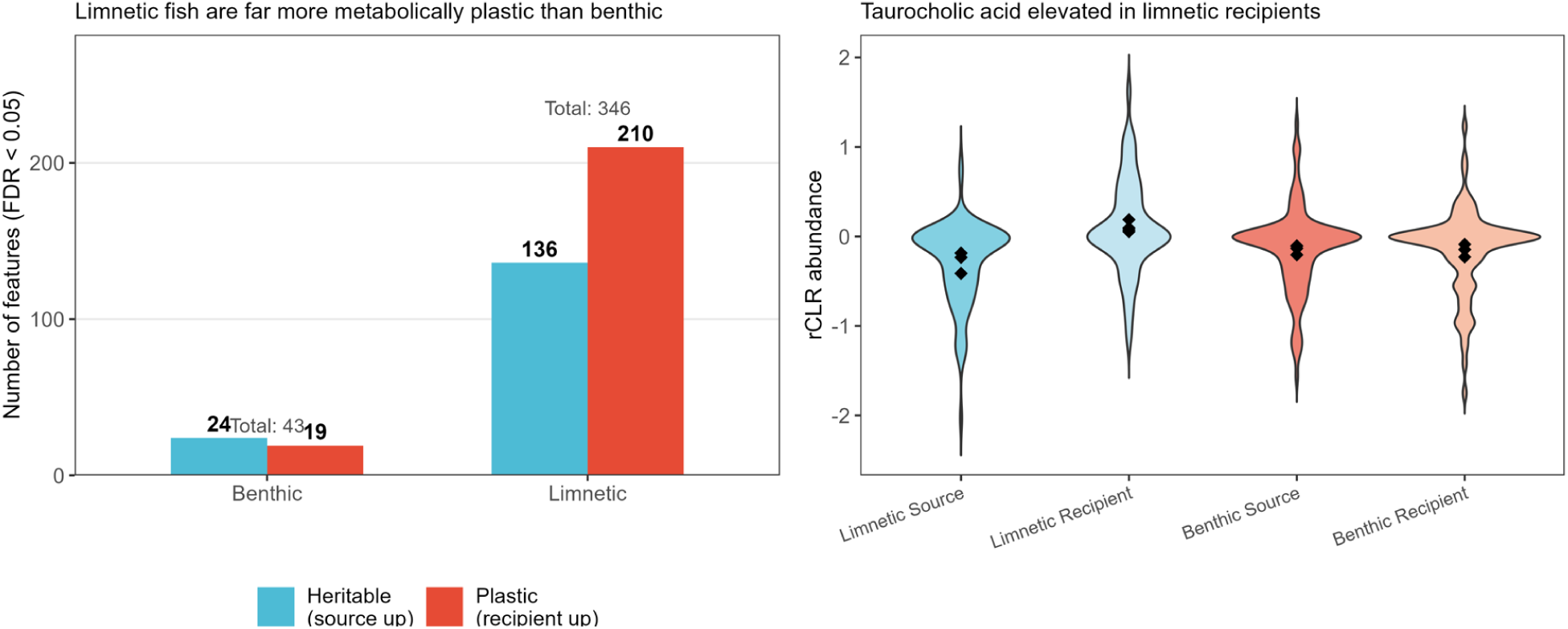
Potential plastic and heritable metabolomic features following translocation, and taurocholic acid elevation in limnetic recipients. A) Number of FDR-significant features (FDR<0.05) classified as potentially heritable (elevated in source fish) or plastic (elevated in recipient fish) in benthic and limnetic fish. B) Taurocholic acid is elevated in limnetic recipients shown through a violin plot with diamonds as the lake means.

Among the three aromatic amino acids consistently elevated in source fish relative to recipient fish across both ecotypes were L-Trytophan (logFC=-0.68, FDR=0.0015), L-Tyrosine (logFC=-0.53, FDR=0.0005), and L-Phenylalanine (logFC=-0.39, FDR=0.05). This differential abundance suggests a potential disruption caused by a novel habitat rather than ancestry where similar quantitative values would be expected from limnetic recipients. These each received MSI Level 2 annotations through GNPS spectral library matching (Figure 5B). All three were also elevated in limnetic source fish relative to benthic source fish (FDR<0.05).

Taurocholic acid (feature 32963, M+Na adduct) was significantly elevated in limnetic ancestry recipients relative to limnetic source fish (logFC=+0.38, FDR=0.013) (Figure 9B;Supplementary Figure S9). Lake habitat type was ruled out as a driver, as all taurocholic acid features were non-significant within limnetic recipients across lake sizes. Taurocholic acid features showed elevated mean abundance in limnetic ancestry groups relative to benthic ancestry groups across both lake sizes, providing a class-level visualization of this pattern (Supplementary Figure S10).

## DISCUSSION

Untargeted metabolomics provides a potentially valuable toolkit for high-dimensional molecular phenotyping of non-model organisms. Fields like ecotoxicology, crop sciences, and animal sciences have been fast to adopt this technology. For instance, LC-MS/MS has been used to study pharmacological effects of diclofenac (DFC, an anti-inflammatory drug) on the stickleback liver metabolome (Lebeau-Roche et al. 2023), environmental effects of altitude on strawberry flavonoid profiles (Rao et al. 2025), and improvements in poultry breeding and meat production (Zhang et al 2024). Untargeted LC-MS/MS is still rarely used in evolutionary biology. There are a growing number of studies comparing metabolomes across different species to document macroevolutionary diversification (Sedio, 2017; Xie et al. 2026), especially in plants (Sedio et al. 2018; Li & Gacquerel, 2021; Schrieber et al. 2023). However, relatively few studies have examined intraspecific metabolomic variation among populations in nature (see Henry et al. 2021; Deng et al. 2025; Nomoto et al. Ramanathan et al. 2026). LC-MS/MS offers a valuable new tool to study microevolutionary diversification within species, phenotypic plasticity, and the physiological basis of adaptation revealed by trait-environment correlations.

### Variation among natural populations of lake stickleback

Our results demonstrate that the liver metabolome differs among closely related lake populations of threespine stickleback. These populations in Cook Inlet, Alaska, have been diverging for approximately 15,000 years since glacial retreat enabled marine fish to establish permanent populations in these isolated freshwater lakes. The populations subsequently diverged genetically, morphologically, and ecologically (Hendry et al. 2024; Bolnick et al. 2026), along a familiar benthic-limnetic ecotype continuum (Willacker et al. 2010; Schluter & MacPhail, 1992). Metabolomic variation allows us to confidently assign individuals to their native population, despite substantial within-lake variation contributed by sexual dimorphism and body size. This result is consistent with studies in other species (mostly plants) finding geographic variation in metabolome composition (Iwanycki et al. 2018; Wang et al. 2025).

The among-population differences in metabolomes can be attributed to both geographic variation (separating the lakes in the Mat-Su versus Kenai regions) and population ecotype (Figure 4; Hendry et al. 2024; Haines et al 2023). Ecotypic divergence was more readily detected by supervised classification (DAPC), but not by unsupervised PCA. This methodological contrast suggests that ecotypic divergence in the metabolome is not concentrated along the highest-variance axes but is instead embedded in lower-variance dimensions that supervised methods are better at isolating. This pattern mirrors findings from genomic studies of stickleback, where ecotype-associated divergence is distributed across many loci of small effect rather than a few large-effect regions (Reid et al. 2021).

Among specific metabolites, limnetic fish showed elevation of metabolites associated with their zooplankton-rich diet relative to benthic fish, which may reflect a generalizable difference in metabolism arising from the divergent diets of these populations. First, limnetics had elevated acylcarnitines, which are intermediates in the process of conjugating long-chain fatty acids to carnitine. Acylcarnitines are also involved in detoxification of acyl groups, regulating fatty acid transport across mitochondrial membranes, and antioxidant functions to reduce oxidative stress. Limnetic ecotype fish might have higher rates of fatty acid catabolism because their zooplankton-diet contains high levels of long-chain polyunsaturated fatty acids (Bjørndal et al. 2018; Hudson et al. 2022). Second, the aromatic amino acids tryptophan, phenylalanine, and tyrosine were significantly elevated in limnetic ecotype fish relative to benthic ecotype fish (FDR<0.05), consistent with zooplankton-rich limnetic diets likely providing higher numbers of aromatic amino acids (Hudson et al. 2022). Vertebrates cannot synthesize these amino acids de novo and must obtain them via their diet or gut microbiome, such as through the shikimate pathway (Parthasarathy et al. 2018; Shende et al. 2024).

### Variation among experimentally founded populations

Comparative studies of phenotypic variation among populations and across environments are beset by a major limitation: genotype and environment are often confounded. Each population may be genetically and ecologically distinct, making it difficult to determine whether phenotype variation is heritable and evolved, or plastic. The experimental translocation of stickleback into fishless lakes allowed us to generate novel permutations of ancestry and environment, by placing fish with benthic-lake ancestry into both small benthic and large limnetic lake habitats (and the converse). We find that habitat, ancestry, and their interaction all contribute to variation in the stickleback liver metabolome (Figure 7B). Metabolomes differed between populations depending on which source lakes were used to establish the population four years (and several generations) earlier. This ancestry effect strongly suggests a heritable component to population differences in the metabolome; some of this represents additive genetic variation, with intermediate discriminant axis scores for mixed-ancestry lakes (Supplementary Figure S7). The habitat effect might arise from any of a number of causes (differences in parasitism, microbiomes, water chemistry, etc), but diet is an obvious candidate. Shifts in diet can alter the liver metabolome within weeks (Hesse et al. 2022), and these lakes harbor distinct communities of prey. Overall, ancestry explained slightly more variation than contemporary habitat, though this is sensitive to the analytical approach (unsupervised versus supervised ordination).

Acylcarnitines, aromatic amino acids, and taurocholic acid connect these population-level patterns to ecotype-specific physiology and potential dietary or microbiome disruption in the novel lake environment. Taurocholic acid elevation in limnetic ancestry recipients was validated through multiple approaches in this study. Taurocholic acid is the primary conjugated bile acid in teleost fish liver, functioning in dietary lipid absorption and cholesterol homeostasis (Xu et al. 2020; Xu et al. 2022). In vertebrates, bile acid conjugation varies with diet, with diets high in animal protein favoring taurine conjugation rather than glycine conjugation (Ridlon et al. 2016). The high animal protein content of zooplankton prey in limnetic source lakes may prime limnetic fish with a better ability to conjugate taurine. Elevated taurocholic acid has been associated with reducing lipid accumulation in the liver (Xu et al. 2022), as well as anti-inflammatory effects in zebrafish (Ge et al. 2023). This concerns limnetic stickleback since they are adapted to zooplankton diets higher in fatty acids and need to avoid conditions like fatty liver disease (Hudson et al. 2022), suggesting dietary fatty acid profiles differ between ecotypes of the native source lakes. Further research could assess whether limnetic populations have locally adapted physiology to this diet high in fatty acids.

Importantly, we also see evidence that habitat matching matters. Phenotypically (and genetically) benthic founder fish placed in limnetic habitats (and limnetic founders in benthic habitats) exhibit some metabolomic signs of convergence towards their new ecotype (implying adaptive plasticity, eg. Figure 8B). However, these mismatched fish also exhibit some distinctive metabolomic profiles that separate them from ecotypes introduced into their pre-adapted habitats (Figure 8A,C). These signals of mismatch might be molecular markers of stress and maladaptation.

This last observation raises an important question: are the metabolomic differences among populations (and ecotypes) adaptive? It is common in comparative studies (within and among species) to use trait-environment correlations to infer adaptive value for traits, a framework established by classic methodological guides (Blanquart et al. 2013; MacColl & Chapman, 2020) and demonstrated in recent empirical gradient studies (Tudor et al. 2024). Following this tradition, we might infer that there is an adaptive basis for ecotype differences in metabolites; though the adaptive divergence might be plastic or environmental. We might go a step further and infer that adaptive evolution produced the inherited ecotype differences separating ancestry treatments in the recipient lakes. However, we should approach such comparative method inferences with particular caution when considering metabolomic features. For example, stickleback exhibit additive genetic variation in morphology and diet (Arnegard et al. 2014). This diet variation is known to impact the stickleback gut microbiome (Bolnick et al. 2014), which in turn produces small molecule metabolites that can enter the host body (Quinn et al. 2020). Thus, differences in metabolite concentrations might be a downstream pleiotropic consequence of heritable variation in ecomorphology and diet. Moreover, higher levels of a particular metabolite in a certain environment are not necessarily indicative of adaptation, but might instead reflect stresses such as immune responses. Such stress responses can be heritable, for instance because genetic variation in diet affects parasite infections (Bolnick et al. 2020). So, although we observe some repeatable ecotype differences in the metabolome, and evidence these are in part heritable and thus a result of evolution, we do not suggest here that these population differences represent local adaptation.

### Limitations of untargeted analyses and chemical annotation

When interpreting untargeted LC-MS/MS data, we must keep some caveats in mind. First and foremost, there is potential for chemical features to be missed. This is especially true for volatile and unstable compounds that do not preserve well in ethanol. For logistical reasons, we were constrained to use ethanol to preserve liver samples in the field, although flash-freezing and long term storage at -80 is preferable to obtain a complete metabolome. Depending on the particular conditions for extraction and LC-MS/MS, certain features can be missed as well. For example, proteomics offers a complementary approach that would detect a different set of (generally larger) molecules that obviously also represent an important set of phenotypes. Second, we only study liver here and are not capturing metabolomic variation in other organs. Each liver was sectioned to separately preserve a portion for metabolomics, and a portion for transcriptomics (subject of a future paper), and there may be variation introduced to our data by the particular sections of liver retained for a given fish (the liver is not a wholly homogenous organ).

Feature annotations are largely putative (MSI Level 2 to 3), and the biological interpretation of our results requires further empirical validation (Rattray et al. 2018). There were also many metabolites that we did not identify precisely, because a majority of chemical compounds detected through LC-MS remain unannotated in databases (da Silva et al. 2015). Consequently, potentially important metabolome variation remains a biological ‘black box’. Future analyses would benefit from using further expanded spectral libraries during the GNPS analysis, integration of dietary and microbiome data, and more controlled laboratory manipulations of genotypes and environment (e.g., diet).

## Conclusions

Untargeted LC-MS/MS can reveal previously undocumented aspects of among-population variation. As with any phenotypes, such variation can arise from both evolved genetic divergence and plastic responses to the environment. In this study, we have provided one of the few empirical studies of among-population metabolomic variation in a wild vertebrate, the threespine stickleback. We show that there are population differences in multiple dimensions of the metabolome. These are driven both by geographic and ecotypic variation, and entail both genetic and plastic effects. For example, benthic versus limnetic ecotypes differ in abundance of acylcarnitines, aromatic amino acids, and taurocholic acid, providing biomarkers for studying the metabolic and molecular basis of (or, consequence of) dietary and ecological variation. These represent candidate molecules for further functional studies, to understand their consequences for organismal performance and fitness. As metabolomic profiling becomes increasingly accessible and cost-effective, we foresee its greater incorporation into studies of local adaptation, ecotypic variation, and microevolutionary change.

## Data Availability

The LC-MS/MS data is available in the public dataset MSV000095708 GNPS - Bolnick Alaska Experimental Evolution (2023) - PRJ1 accessible on the MassIVE repository: https://massive.ucsd.edu/ProteoSAFe/dataset.jsp?task=8b2ff40d4b694adc85c4ea605b1a4622. The processed feature tables and peak lists for GNPS2, and mgf file for SIRIUS annotations are also available in the public dataset MSV000095708 on MassIVE. The R scripts and data tables to replicate our analyses and graphics are publicly archived as a Figshare repository at doi:10.6084/m9.figshare.32745483.

## Author contributions

Conceptualization: DIB, PCD, WHFD, MG. Data Curation: AMC-R, MMT, MG, WHFD, DIB. Formal analysis: WHFD, MG, DIB, AMC-R. Funding acquisition: DIB, PCD, CLP. Methodology: PCD, AMC-R, DIB, APH, CLP, RDHB. Investigation: WHFD, MG, MMT, AMC-R, AL, MW, Y-JT. Project administration: DIB, PCD, AL, CLP. Resources: DIB, CLP, APH, AL, KMM, JNW, NCS. Supervision. DIB, PCD. Visualization: WHFD, MG, DIB. Writing: WHFD, MG, MMT, DIB, AMC-R. Reviewing & editing: all authors.

## Acknowledgements

We thank the Alaska Department of Fish and Game, and the Kenai National Wildlife Refuge, and Alaska State Parks for cooperation in permitting. We thank local landowners for access to the research sites through their private property.

## Supplementary

**Supplementary Figure S1.**
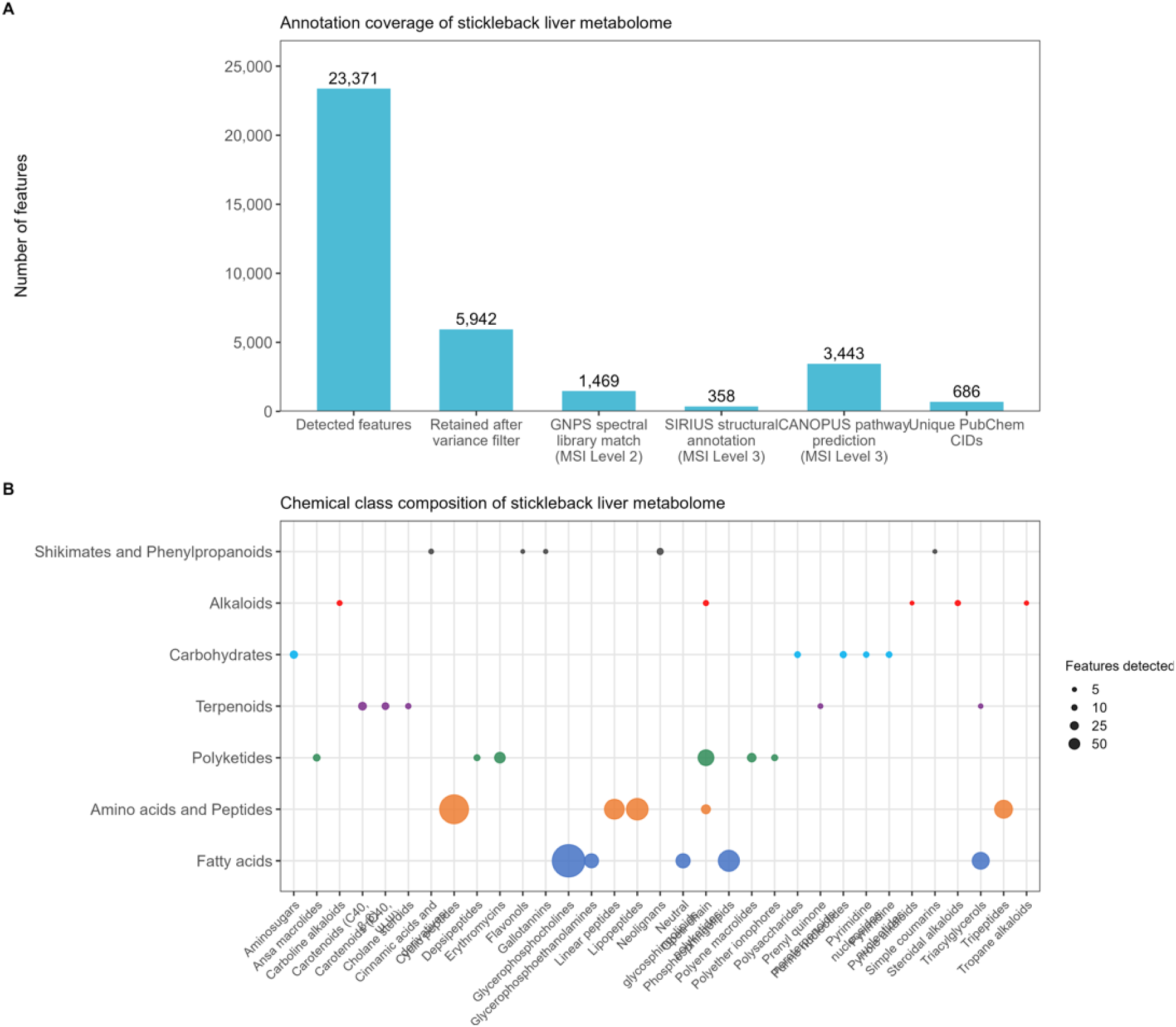
Annotation coverage and chemical class composition of the stickleback liver metabolome. Panel A: Number of features at each stage of detection and annotation across the three-tier MSI annotation pipeline. Panel B: Bubble size reflects the total abundance of features across samples per chemical class predicted by CANOPUS NPClassifier (n=3,443) features with class-level predictions, MS-222 and DEET excluded). Rows represent a broad NPClassifier class, and columns represent a chemical subclass within that class but are limited to the top five most abundant subclasses. All annotations here are putative (MSI Level 3).

**Supplementary Figure S2.**
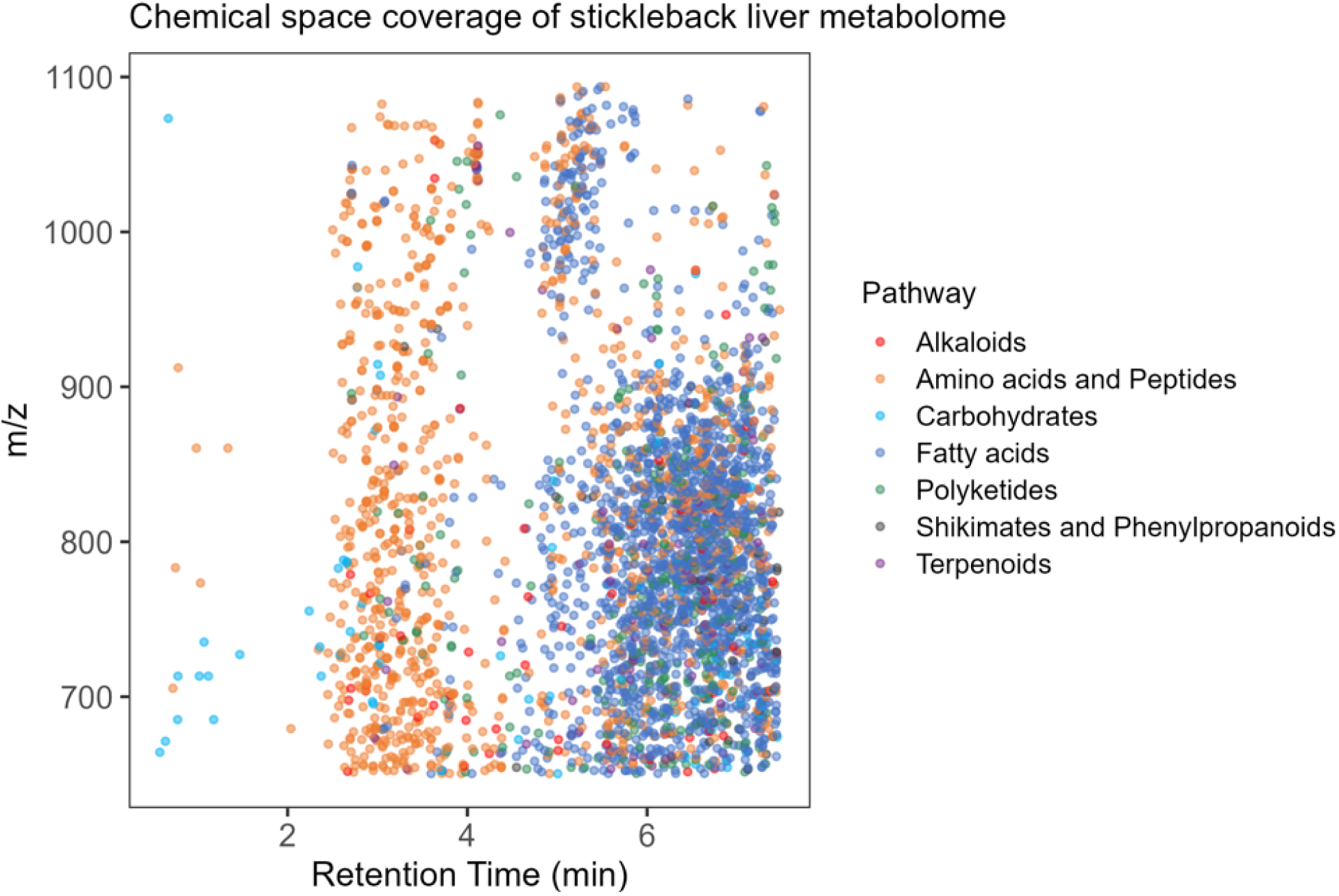
Retention time versus m/z ratio showing chemical space of CANOPUS-annotated features colored by NPClassifier class as they elute from the reverse-phase C18 LC column.

**Supplementary Figure S3.**
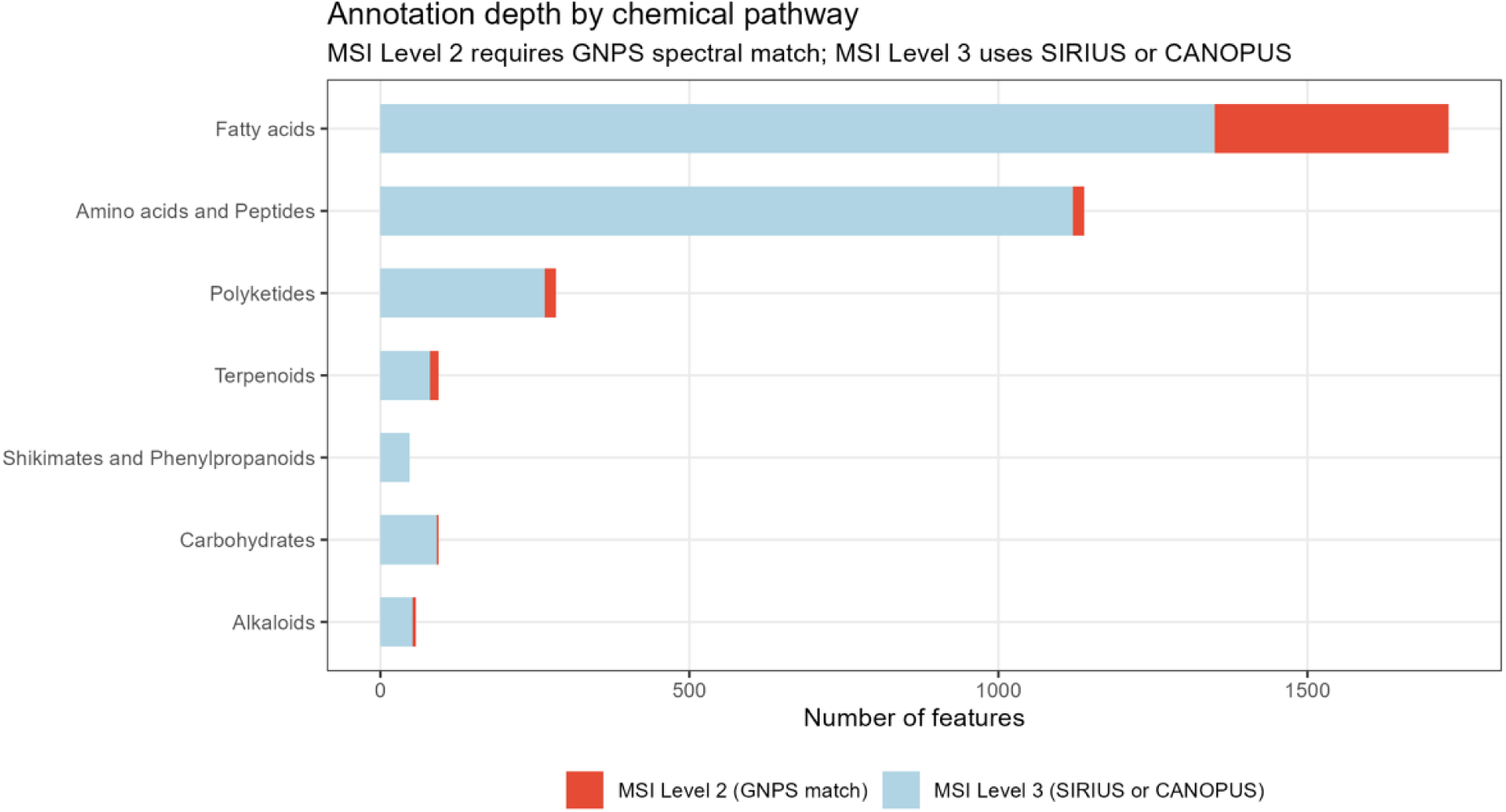
Annotation depth by chemical class. Stacked bars show the number of features per class annotated at MSI Level 2 compared to MSI Level 3.

**Supplementary Figure S4.**
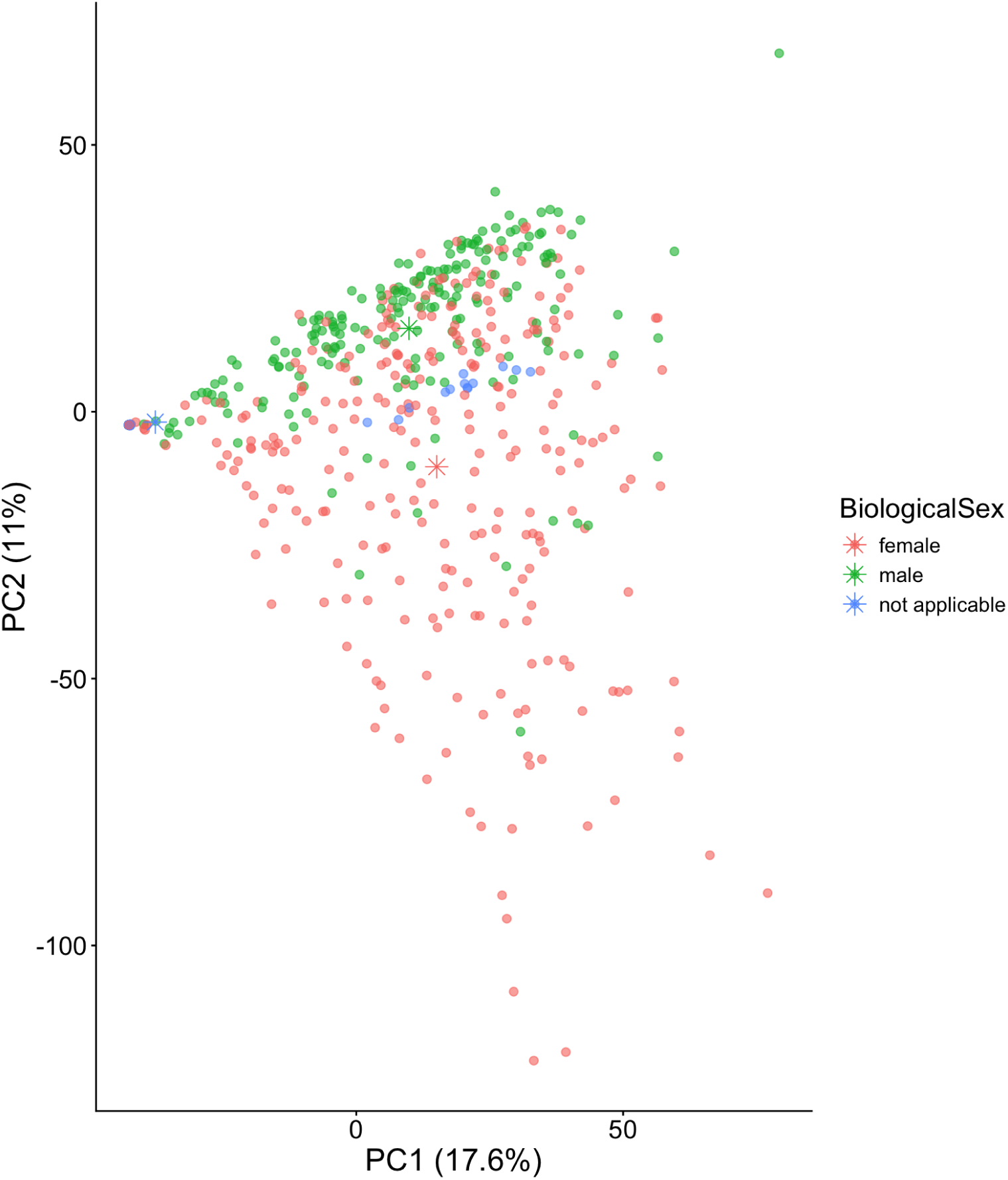
Sexual dimorphism in stickleback metabolomes for the full dataset of 480 individuals. Stickleback with undetermined sex (by gonad morphology) are listed as “not applicable”.

**Supplementary Figure S5.**
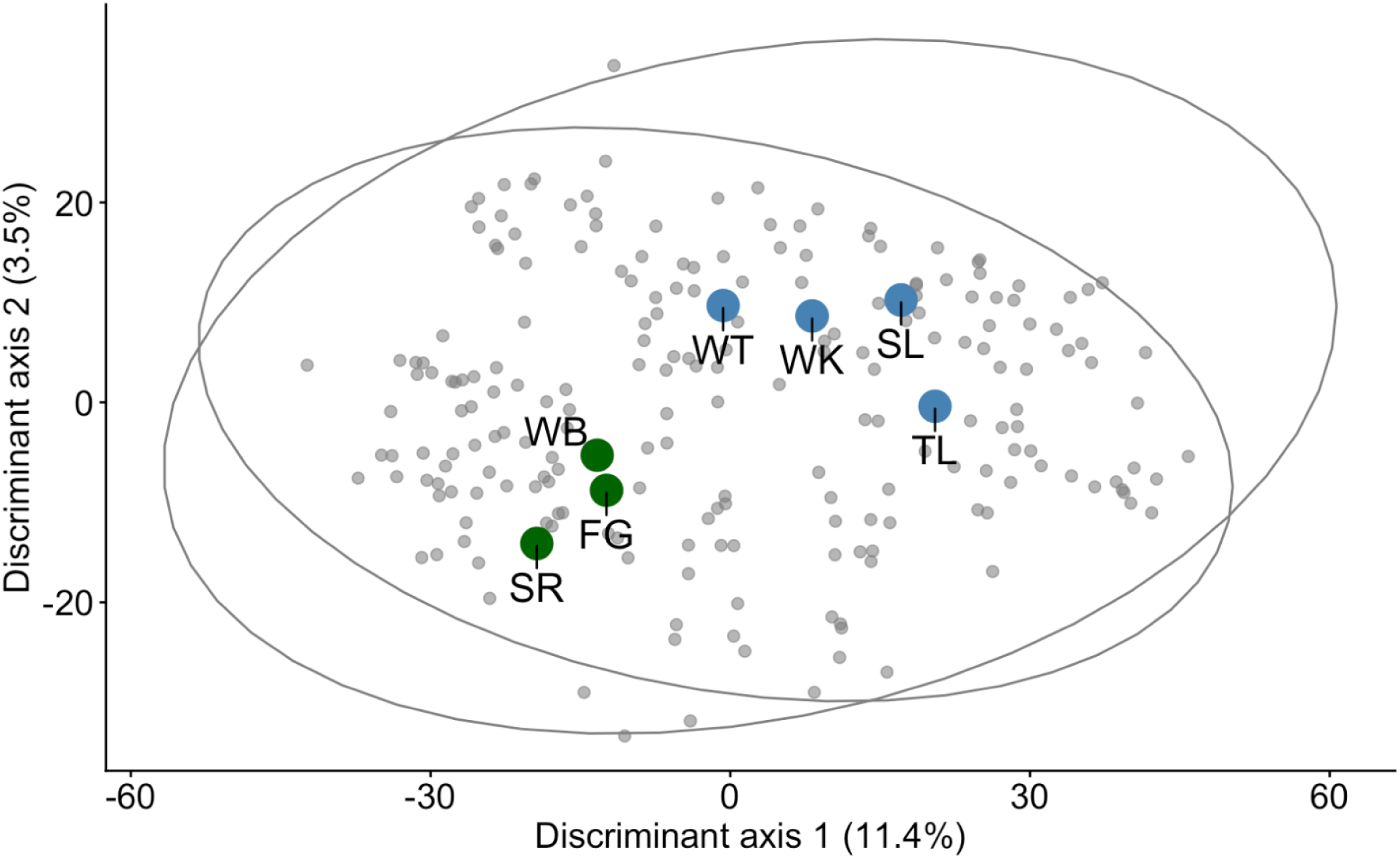
PLSDA supervised classification of Kenai Peninsula (blue) versus Matsu Valley (green) source lake populations.

**Supplementary Figure S6:**
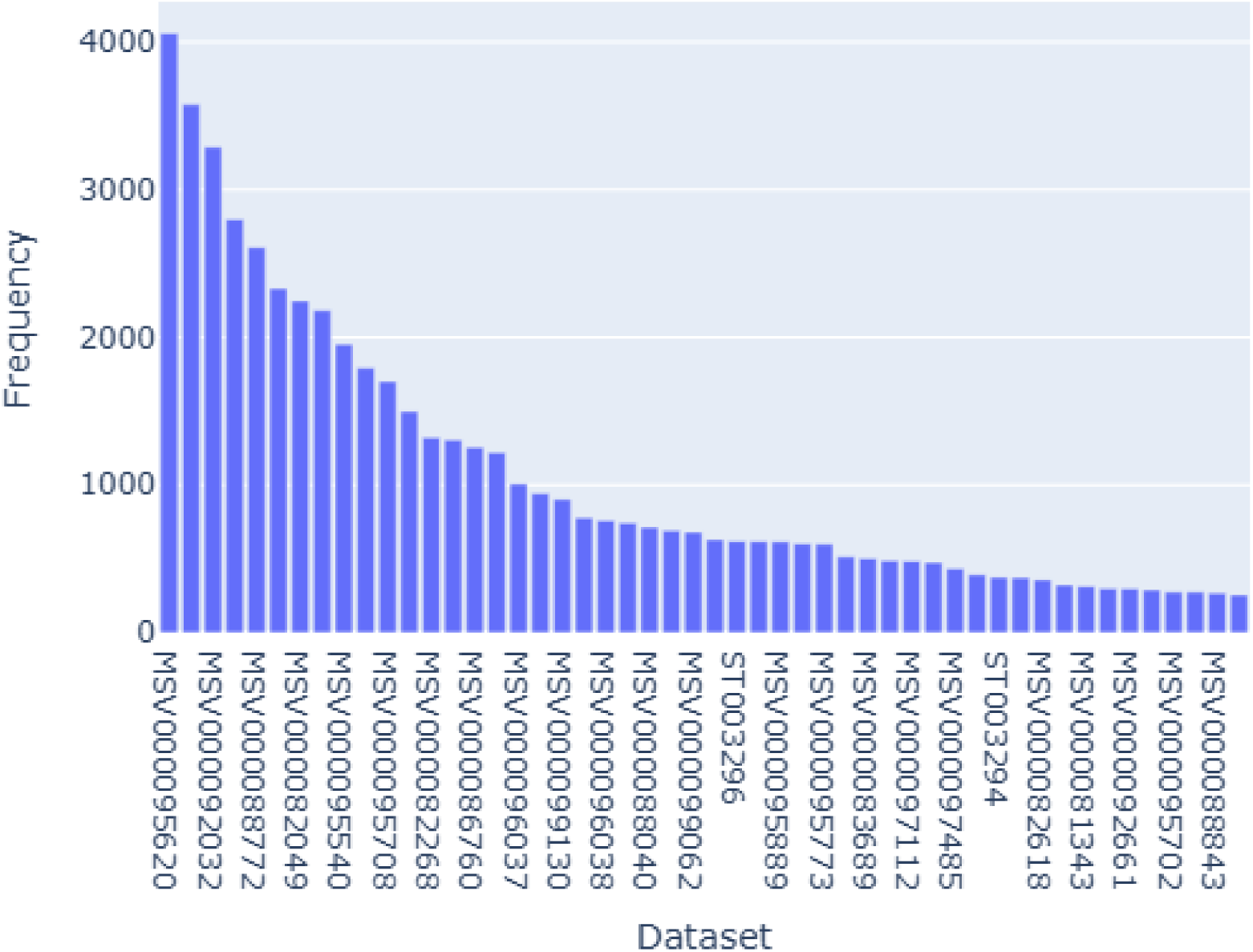
Datasets with the highest frequencies of taurocholic acid feature 32963. http://fasst.gnps2.org/fastsearch/?usi1=mzspec%3AGNPS2%3ATASK-a78a3532e6f14df38f61183b0835ed53-nf_output%2Fclustering%2Fspectra_reformatted.mgf%3Ascan%3A32963&precursor_mz=None&charge=None&library_select=metabolomicspanrepo_index_nightly&analog_select=No&delta_mass_below=130&delta_mass_above=200&pm_tolerance=0.05&fragment_tolerance=0.05&cosine_threshold=0.7&use_peaks=0#%7B%22peaks%22%3A%20null%7D

**Supplementary Figure S7.**
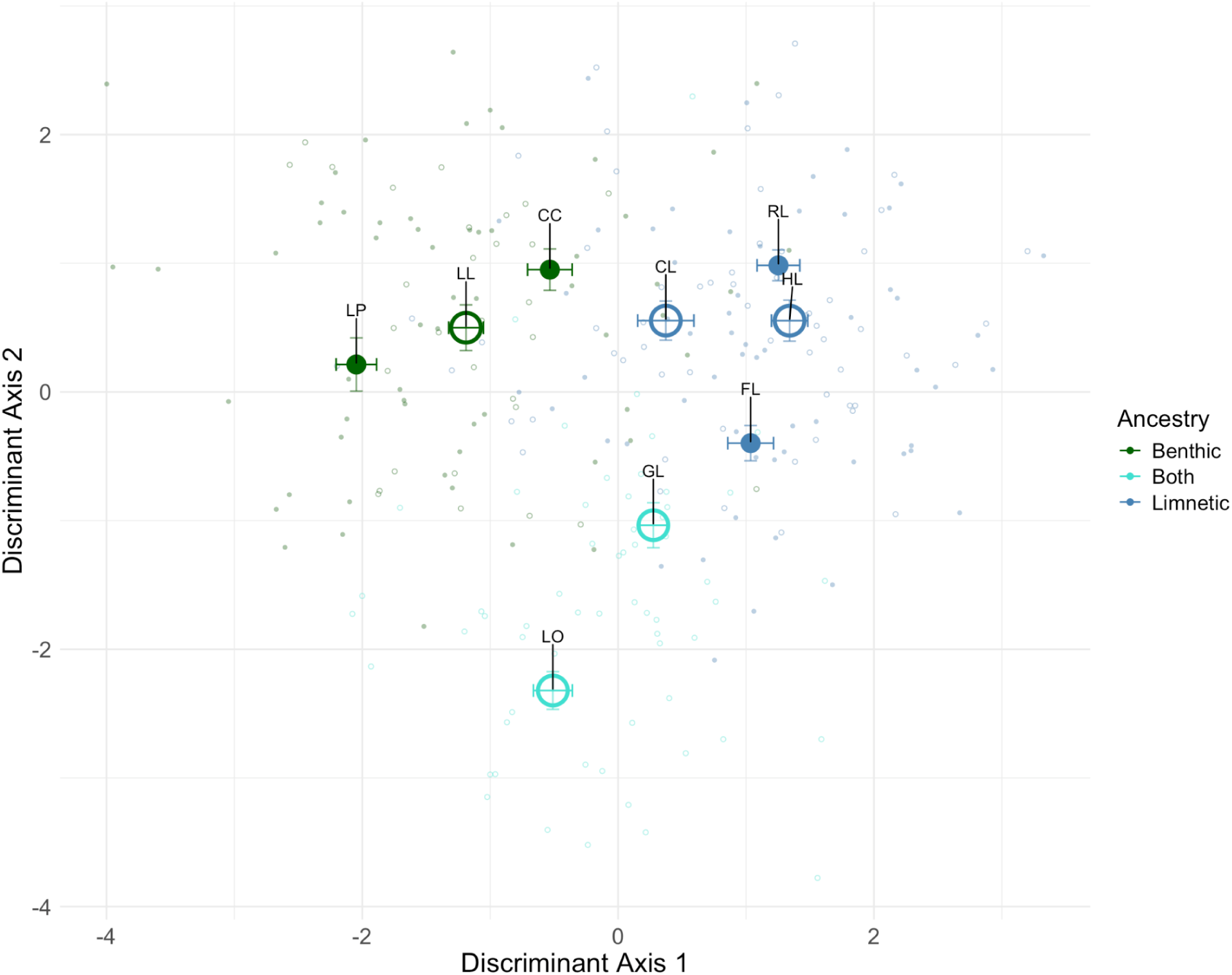
A DAPC supervised classification of recipient lake fish, trained only on ecotype (benthic ancestry or limnetic ancestry) places the mixed-ancestry lakes (G Lake, GL; Loon Lake, LO) in the center of the benthic-limnetic continuum, implying some additive genetic control of the benthic-limnetic variation in the metabolome.

**Supplementary Figure S8.**
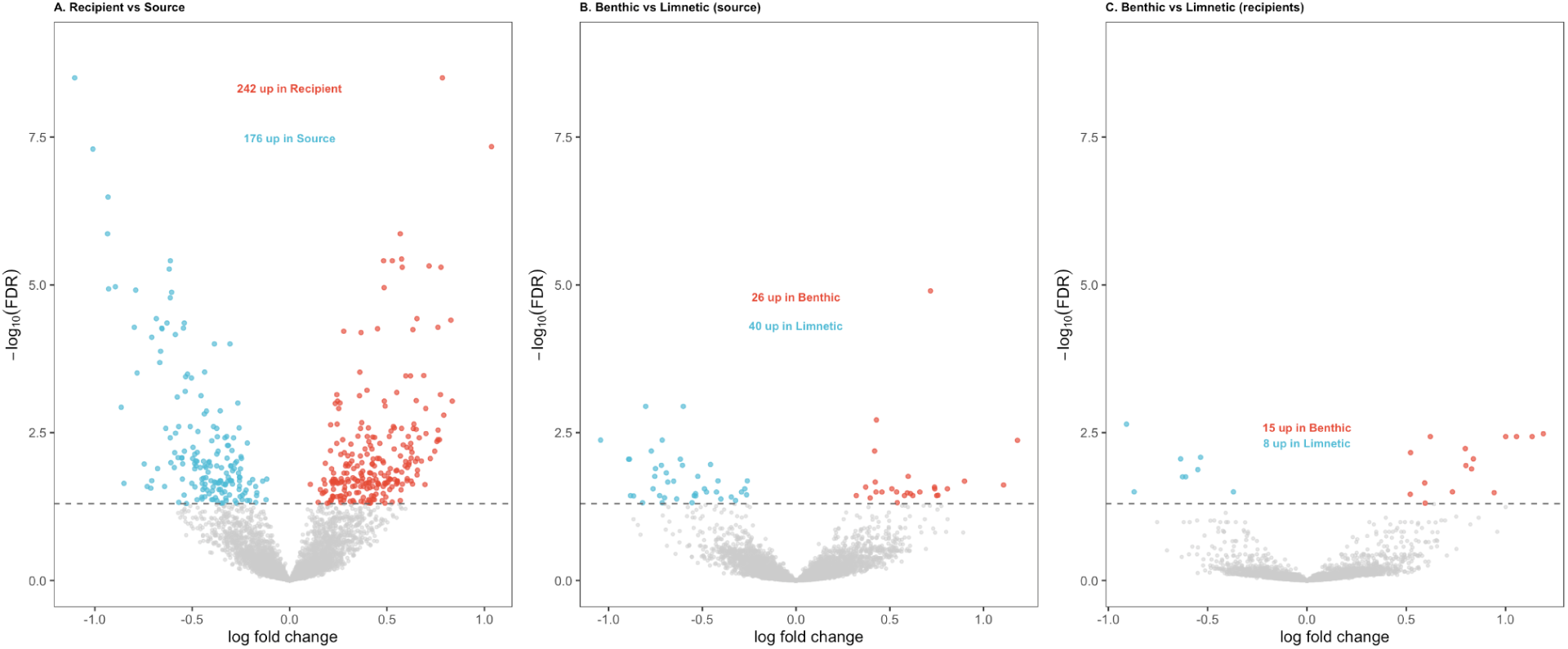
Volcano plots showing metabolome feature differential abundance between A) source versus recipient lakes, B) benthic versus limnetic ecotype source lakes, and C) benthic- versus limnetic-ancestry recipient lakes. The x axis is log fold change, a measure of effect size and direction; red points are metabolites which are more abundant in recipient (A) or benthic (B,C) fish, blue are more abundant in source (or limnetic) fish. The y axis is the log P-value measuring strength of statistical confidence in rejecting the null hypothesis.

**Supplementary Figure S9.**
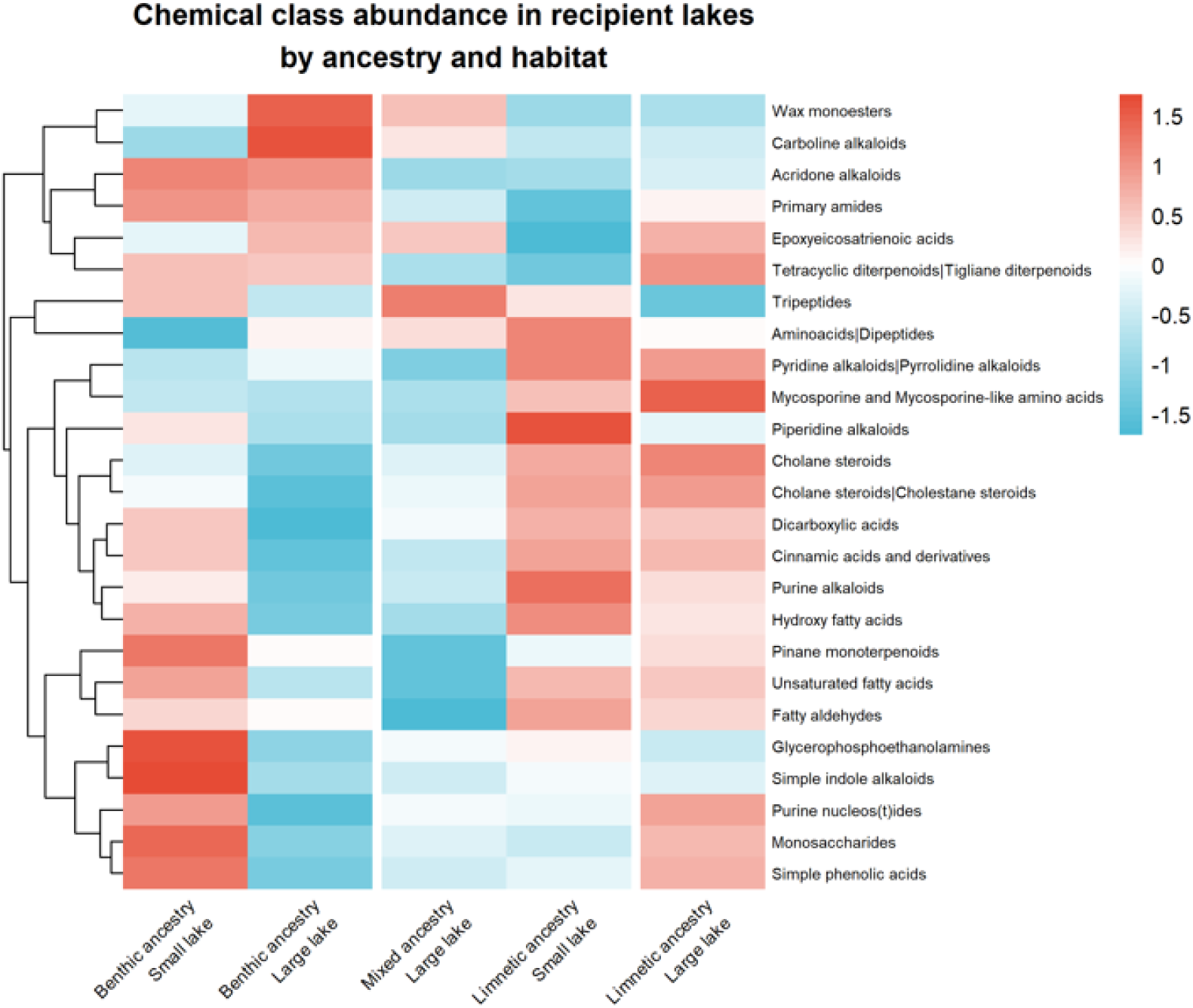
Chemical class abundance in recipient lakes by ancestry and habitat. Heatmap shows the top 25 chemical classes by variance across groups, scaled by row (z-score). Columns represent lake groups organized by ancestral genotype pool and lake size. Rows are clustered by hierarchical clustering. Red indicates elevated abundance relative to the row mean; blue indicates reduced abundance.

**Supplementary Figure S10.**
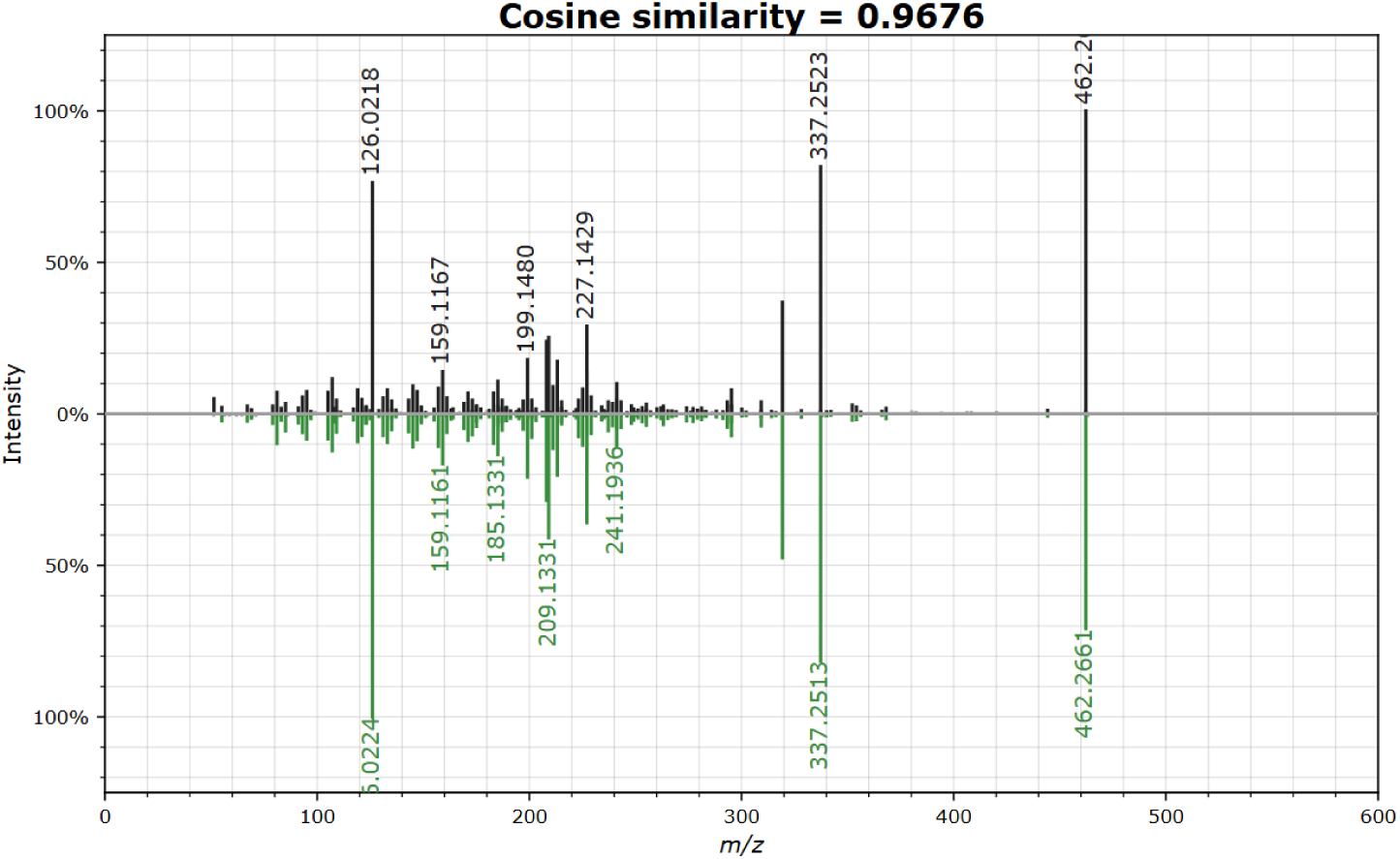
Mirror spectrum confirming taurocholic acid annotation for feature 32958. Black peaks show the detected spectrum; green peaks show the library reference spectrum (Library ID: CCMSLIB00005435564). Feature 32958 shares retention time (4.12 min) with feature 32963 and both cluster in the same molecular family on Cytoscape. Feature 32963 co-occurs in the same molecular family as feature 32958 (M-3H2O+H adduct, cosine=0.970), which provides spectral annotation for taurocholic acid. Both features share identical retention time (4.12 min) and cluster together in the FBMN molecular network (Figure 6A), confirming they represent different forms of the same compound (). This equivalence between these molecular features is supported by: 1) differential abundance analysis (logFC=+0.38, FDR=0.013); 2) MSI Level 2 GNPS spectral match (cosine similar=0.864 for feature 32963, cosine=0.970 for feature 32958); 3) fastMASST spectral search identifying matches in two of our other public repositories of stickleback metabolomics data: one project observing stickleback liver metabolome data from 47 Vancouver Island lakes (MSV000095784) and another study looking at immune activation and fibrosis (MSV000095620).

**Supplementary Figure S11:**
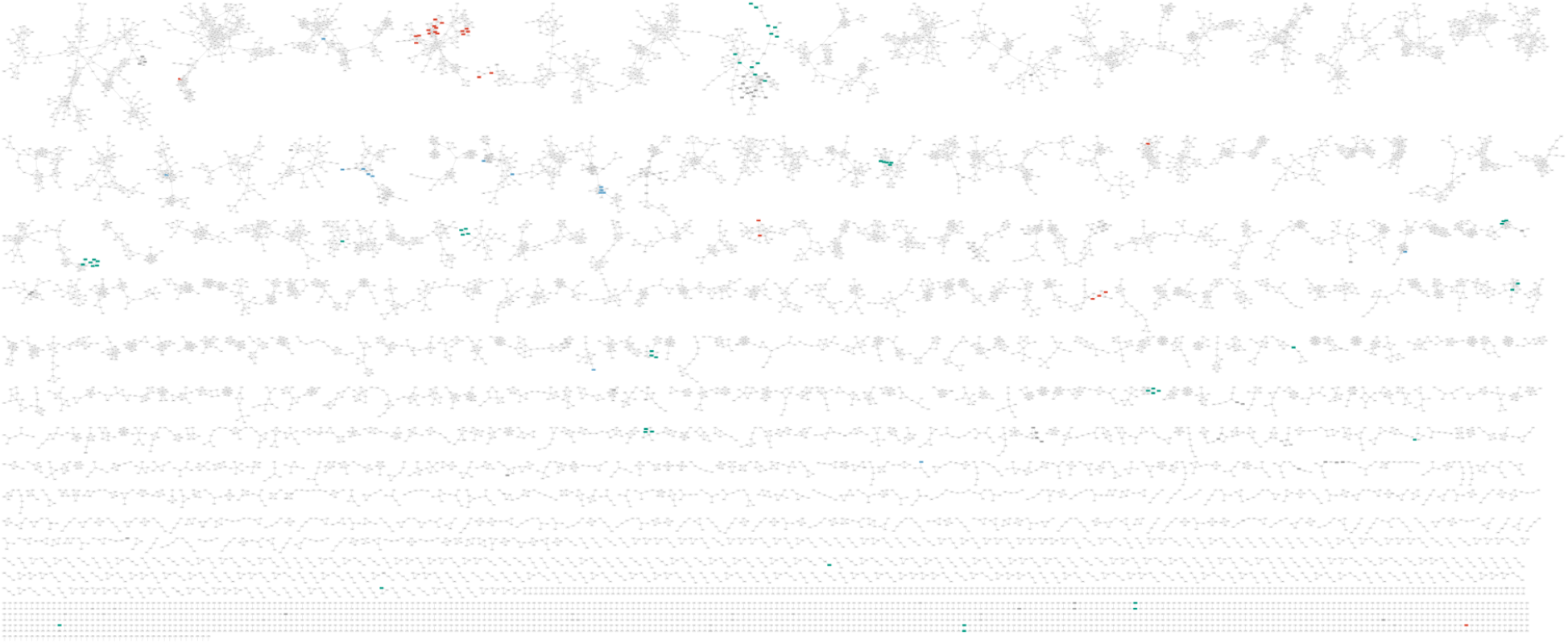
Molecular network of the stickleback liver metabolome visualized in Cytoscape. Each node is an individual feature. Edges represent cosine similarity > or equal to0.7 with > or equal to 4 matched peaks. Node color indicates NPClassifier chemical class assignment: red=cholane steroids, teal=amino acids, blue=lipopeptides, and grey=unannotated and all other classes. All annotations are putative (MSI 2 and MSI 3).

## REFERENCES

Alanazi, S. (2025). Recent Advances in Liquid Chromatography–Mass Spectrometry (LC–MS) Applications in Biological and Applied Sciences. Analytical Science Advances, 6(1), e70024. 10.1002/ansa.70024

Arnegard, M. E., McGee, M. D., Matthews, B., Marchinko, K. B., Conte, G. L., Kabir, S., Bedford, N., Bergek, S., Chan, Y. F., Jones, F. C., Kingsley, D. M., Peichel, C. L., & Schluter, D. (2014). Genetics of ecological divergence during speciation. Nature, 511(7509), 307–311. 10.1038/nature13301

Bell, M. A., & Foster, S. A. (Eds.). (1994). The Evolutionary Biology of the Threespine Stickleback. Oxford University Press Oxford. 10.1093/oso/9780198577287.001.0001

Bittremieux, W., Chen, C., Dorrestein, P. C., Schymanski, E. L., Schulze, T., Neumann, S., Meier, R., Rogers, S., & Wang, M. (2020). Universal MS/MS Visualization and Retrieval with the Metabolomics Spectrum Resolver Web Service. Bioinformatics. 10.1101/2020.05.09.086066

Bjørndal, B., Alterås, E. K., Lindquist, C., Svardal, A., Skorve, J., & Berge, R. K. (2018). Associations between fatty acid oxidation, hepatic mitochondrial function, and plasma acylcarnitine levels in mice. Nutrition & Metabolism, 15(1), 10. 10.1186/s12986-018-0241-7

Blanquart, F., Kaltz, O., Nuismer, S. L., & Gandon, S. (2013). A practical guide to measuring local adaptation. Ecology Letters, 16(9), 1195–1205. 10.1111/ele.12150

Bolnick, D. I., & Ballare, K. M. (2020). Resource diversity promotes among-individual diet variation, but not genomic diversity, in lake stickleback. Ecology Letters, 23(3), 495–505. 10.1111/ele.13448

Bolnick, D. I., Eckert, L., Barrett, R. D. H., Choi, E., Haines, G., Hendry, A. P., Kerns, E. V., Lind, Å. J., Milligan, -McClellan Kathryn, Peichel, C. L., Sasser, K., Thornton, A. R., Wolf, C., Steinel, N. C., & Weber, J. N. (n.d.). Eco-evolutionary dynamics of infection and immunity in experimentally founded lake populations of threespine stickleback. The American Naturalist, 0(ja). 10.1086/741682

Bolnick, D. I., Resetarits, E. J., Ballare, K., Stuart, Y. E., & Stutz, W. E. (2020). Scale-dependent effects of host patch traits on species composition in a stickleback parasite metacommunity. Ecology, 101(12), e03181. 10.1002/ecy.3181

Bolnick, D. I., Snowberg, L. K., Hirsch, P. E., Lauber, C. L., Org, E., Parks, B., Lusis, A. J., Knight, R., Caporaso, J. G., & Svanbäck, R. (2014). Individual diet has sex-dependent effects on vertebrate gut microbiota. Nature Communications, 5(1), 4500. 10.1038/ncomms5500

Brachmann, M. K., Smith, B., Kristjánsson, B., Selman, C., & Parsons, K. (2026). Energetic misfires: Hybridization drives transgressive expression in metabolic pathways in thermally divergent Icelandic stickleback (p. 2026.04.17.719192). bioRxiv. 10.64898/2026.04.17.719192

Cao, M., Fraser, K., Jones, C., Stewart, A., Lyons, T., Faville, M., & Barrett, B. (2017). Untargeted Metabotyping Lolium perenne Reveals Population-Level Variation in Plant Flavonoids and Alkaloids. Frontiers in Plant Science, 8. 10.3389/fpls.2017.00133

Caraballo, M., Tawfik, M., Tseng, J., & Wu, M. (2024). Extraction Protocol for untargeted LC-MS/MS - Animal tissues. protocols.io 10.17504/protocols.io.6qpvr8x8blmk/v1

Chen, H., Kong, J., Du, P., Wang, Q., Jiang, T., Hou, X., Feng, T., Duan, J., & Liu, C. (2025). Functional metabolomics: Unlocking the role of small molecular metabolites. Frontiers in Molecular Biosciences, 12. 10.3389/fmolb.2025.1542100

Clish, C. B. (2015). Metabolomics: An emerging but powerful tool for precision medicine. Molecular Case Studies, 1(1), a000588. 10.1101/mcs.a000588

Coler, E. A., Melnik, A., Lotfi, A., Moradi, D., Ahiadu, B., Portal Gomes, P. W., Patan, A., Charron-Lamoureux, V., Dorrestein, P. C., Barnes, S., Boginski, V., Semenov, A., & Aksenov, A. A. (2026). Ordering molecular diversity in untargeted metabolomics via molecular community networking. Cell Reports Methods, 101468. 10.1016/j.crmeth.2026.101468

da Silva, R. R., Dorrestein, P. C., & Quinn, R. A. (2015). Illuminating the dark matter in metabolomics. Proceedings of the National Academy of Sciences, 112(41), 12549–12550. 10.1073/pnas.1516878112

Deng, L.-J., Li, Y.-L., Wang, F.-Y., Sun, X.-Q., Milne, R. I., Liu, J., & Wu, Z.-Y. (2025). Comparative metabolomics of two nettle species unveils distinct high-altitude adaptation mechanisms on the Tibetan Plateau. BMC Plant Biology, 25(1), 640. 10.1186/s12870-025-06666-9

Domer, A., Jasinska, W., Rosental, L., Shochat, E., Alseekh, S., Fernie, A. R., Brotman, Y., & Ovadia, O. (2025). Comparative analysis of the plasma metabolome of migrating passerines: Novel insights into stopover metabolism. Journal of Avian Biology, 2025(2), e03331. 10.1111/jav.03331

Dührkop, K., Fleischauer, M., Ludwig, M., Aksenov, A. A., Melnik, A. V., Meusel, M., Dorrestein, P. C., Rousu, J., & Böcker, S. (2019). SIRIUS 4: A rapid tool for turning tandem mass spectra into metabolite structure information. Nature Methods, 16(4), 299–302. 10.1038/s41592-019-0344-8

Dührkop, K., Nothias, L.-F., Fleischauer, M., Reher, R., Ludwig, M., Hoffmann, M. A., Petras, D., Gerwick, W. H., Rousu, J., Dorrestein, P. C., & Böcker, S. (2021). Systematic classification of unknown metabolites using high-resolution fragmentation mass spectra. Nature Biotechnology, 39(4), 462–471. 10.1038/s41587-020-0740-8

Eckert, L., Bolnick, D. I., Derry, A. M., Haines, G. E., Heckley, A. M., Lind, Å. J., Peichel, C. L., Roth, A. M., Steinel, N. C., Vlahiotis, K., Weber, J. N., Hendry, A. P., & Barrett, R. D. H. (2026). Intrinsic fitness differences outweigh environmental matching in shaping introduction outcomes in nature (p. 2026.02.04.699496). bioRxiv. 10.64898/2026.02.04.699496

Elser, D., Pflieger, D., Villette, C., Moegle, B., Miesch, L., & Gaquerel, E. (2023). Evolutionary metabolomics of specialized metabolism diversification in the genus Nicotiana highlights N-acylnornicotine innovations. Science Advances, 9(34), eade8984. 10.1126/sciadv.ade8984

Ge, X., Huang, S., Ren, C., & Zhao, L. (2023). Taurocholic Acid and Glycocholic Acid Inhibit Inflammation and Activate Farnesoid X Receptor Expression in LPS-Stimulated Zebrafish and Macrophages. Molecules, 28(5), 2005. 10.3390/molecules28052005

Giebułtowicz, J., Grabicová, K., Brooks, B. W., & Grabic, R. (2024). Influence of time-dependent sampling on the plasma metabolome and exposome of fish collected from an effluent-dependent pond. Science of The Total Environment, 906, 167446. 10.1016/j.scitotenv.2023.167446

Haines, G. E., Moisan, L., Derry, A. M., & Hendry, A. P. (2023). Dimensionality and Modularity of Adaptive Variation: Divergence in Threespine Stickleback from Diverse Environments. The American Naturalist, 201(2), 175–199. 10.1086/722483

Han, Z., Sun, J., Jiang, B., Chen, K., Ge, L., Sun, Z., & Wang, A. (2024). Fecal microbiota transplantation accelerates restoration of florfenicol-disturbed intestinal microbiota in a fish model. Communications Biology, 7(1), 1006. 10.1038/s42003-024-06727-z

Hendry, A. P., Barrett, R. D. H., Bell, A. M., Bell, M. A., Bolnick, D. I., Gotanda, K. M., Haines, G. E., Lind, Å. J., Packer, M., Peichel, C. L., Peterson, C. R., Poore, H. A., Massengill, R. L., Milligan-McClellan, K., Steinel, N. C., Sanderson, S., Walsh, M. R., Weber, J. N., & Derry, A. M. (2024). Designing eco-evolutionary experiments for restoration projects: Opportunities and constraints revealed during stickleback introductions. Ecology and Evolution, 14(6), e11503. 10.1002/ece3.11503

Henry, J. A., Khattri, R. B., Guingab-Cagmat, J., Merritt, M. E., Garrett, T. J., Patterson, J. T., & Lohr, K. E. (2021). Intraspecific variation in polar and nonpolar metabolite profiles of a threatened Caribbean coral. Metabolomics: Official Journal of the Metabolomic Society, 17(7), 60. 10.1007/s11306-021-01808-0

Hesse, T., Nachev, M., Khaliq, S., Jochmann, M. A., Franke, F., Scharsack, J. P., Kurtz, J., Sures, B., & Schmidt, T. C. (2022). Insights into amino acid fractionation and incorporation by compound-specific carbon isotope analysis of three-spined sticklebacks. Scientific Reports, 12(1), 11690. 10.1038/s41598-022-15704-7

Heuckeroth, S., Damiani, T., Smirnov, A., Mokshyna, O., Brungs, C., Korf, A., Smith, J. D., Stincone, P., Dreolin, N., Nothias, L.-F., Hyötyläinen, T., Orešič, M., Karst, U., Dorrestein, P. C., Petras, D., Du, X., van der Hooft, J. J. J., Schmid, R., & Pluskal, T. (2024). Reproducible mass spectrometry data processing and compound annotation in MZmine 3. Nature Protocols, 19(9), 2597–2641. 10.1038/s41596-024-00996-y

Hong, Y., Chen, B., Zhai, X., Qian, Q., Gui, R., & Jiang, C. (2023). Integrated analysis of the gut microbiome and metabolome in a mouse model of inflammation-induced colorectal tumors. Frontiers in Microbiology, 13. 10.3389/fmicb.2022.1082835

Hudson, C. M., Ladd, S. N., Leal, M. C., Schubert, C. J., Seehausen, O., & Matthews, B. (2022). Fit and fatty freshwater fish: Contrasting polyunsaturated fatty acid phenotypes between hybridizing stickleback lineages. Oikos, 2022(7). 10.1111/oik.08558

Ishikawa, A., Kabeya, N., Ikeya, K., Kakioka, R., Cech, J. N., Osada, N., Leal, M. C., Inoue, J., Kume, M., Toyoda, A., Tezuka, A., Nagano, A. J., Yamasaki, Y. Y., Suzuki, Y., Kokita, T., Takahashi, H., Lucek, K., Marques, D., Takehana, Y., … Kitano, J. (2019). A key metabolic gene for recurrent freshwater colonization and radiation in fishes. Science, 364(6443), 886–889. 10.1126/science.aau5656

Ishikawa, A., Stuart, Y. E., Bolnick, D. I., & Kitano, J. (2021). Copy number variation of a fatty acid desaturase gene Fads2 associated with ecological divergence in freshwater stickleback populations. Biology Letters, 17(8), 20210204. 10.1098/rsbl.2021.0204

Iwanycki Ahlstrand, N., Havskov Reghev, N., Markussen, B., Bruun Hansen, H. C., Eiriksson, F. F., Thorsteinsdóttir, M., Rønsted, N., & Barnes, C. J. (2018). Untargeted metabolic profiling reveals geography as the strongest predictor of metabolic phenotypes of a cosmopolitan weed. Ecology and Evolution, 8(13), 6812–6826. 10.1002/ece3.4195

Jeffries, K. M., Connon, R. E., Davis, B. E., Komoroske, L. M., Britton, M. T., Sommer, T., Todgham, A. E., & Fangue, N. A. (2016). Effects of high temperatures on threatened estuarine fishes during periods of extreme drought. Journal of Experimental Biology, 219(11), 1705–1716. 10.1242/jeb.134528

Jombart, T. (2008). adegenet: A R package for the multivariate analysis of genetic markers. Bioinformatics, 24(11), 1403–1405. 10.1093/bioinformatics/btn129

Jombart, T., & Ahmed, I. (2011). adegenet 1.3-1: New tools for the analysis of genome-wide SNP data. Bioinformatics, 27(21), 3070–3071. 10.1093/bioinformatics/btr521

Jombart, T., Devillard, S., & Balloux, F. (2010). Discriminant analysis of principal components: A new method for the analysis of genetically structured populations. BMC Genetics, 11(1), 94. 10.1186/1471-2156-11-94

Jones, F. C., Grabherr, M. G., Chan, Y. F., Russell, P., Mauceli, E., Johnson, J., Swofford, R., Pirun, M., Zody, M. C., White, S., Birney, E., Searle, S., Schmutz, J., Grimwood, J., Dickson, M. C., Myers, R. M., Miller, C. T., Summers, B. R., Knecht, A. K., … Kingsley, D. M. (2012). The genomic basis of adaptive evolution in threespine sticklebacks. Nature, 484(7392), 55–61. 10.1038/nature10944

Kim, H. W., Wang, M., Leber, C. A., Nothias, L.-F., Reher, R., Kang, K. B., van der Hooft, J. J. J., Dorrestein, P. C., Gerwick, W. H., & Cottrell, G. W. (2021). NPClassifier: A Deep Neural Network-Based Structural Classification Tool for Natural Products. Journal of Natural Products, 84(11), 2795–2807. 10.1021/acs.jnatprod.1c00399

Klausmeier, C. A., Kremer, C. T., & Koffel, T. (2020). Trait-based ecological and eco-evolutionary theory. In C. A. Klausmeier, C. T. Kremer, & T. Koffel, Theoretical Ecology (pp. 161–194). Oxford University Press. 10.1093/oso/9780198824282.003.0011

Kolde, R. (2025). pheatmap: Pretty Heatmaps (Version 1.0.13) [Computer software]. https://cran.r-project.org/web/packages/pheatmap/index.html

Kuhn, M. (2008). Building Predictive Models in R Using the caret Package. Journal of Statistical Software, 28, 1–26. 10.18637/jss.v028.i05

Langfelder, P., & Horvath, S. (2008). WGCNA: An R package for weighted correlation network analysis. BMC Bioinformatics, 9(1), 559. 10.1186/1471-2105-9-559

Lankadurai, B. P., Nagato, E. G., & Simpson, M. J. (2013). Environmental metabolomics: An emerging approach to study organism responses to environmental stressors. Environmental Reviews, 21(3), 180–205. 10.1139/er-2013-0011

Lavin, P. A., & Mcphail, J. D. (1985). The evolution of freshwater diversity in the threespine stickleback (Gasterosteus aculeatus): Site-specific differentiation of trophic morphology. Canadian Journal of Zoology, 63(11), 2632–2638. 10.1139/z85-393

Lavin, P. A., & McPhail, J. D. (1986). Adaptive Divergence of Trophic Phenotype among Freshwater Populations of the Threespine Stickleback (Gasterosteus aculeatus). Canadian Journal of Fisheries and Aquatic Sciences, 43(12), 2455–2463. 10.1139/f86-305

Lebeau-Roche, E., Daniele, G., Fildier, A., Bonnefoy, C., Turiès, C., Bado-Nilles, A., Porcher, J.-M., Dedourge-Geffard, O., Vulliet, E., & Geffard, A. (2023). Time and dose-dependent impairment of liver metabolism in Gasterosteus aculeatus following exposure to diclofenac (DCF) highlighted by LC-HRMS untargeted metabolomics. The Science of the Total Environment, 858(Pt 1), 159801. 10.1016/j.scitotenv.2022.159801

Li, B., Liang, C., Xu, B., Song, P., Liu, D., Zhang, J., Gu, H., Jiang, F., Gao, H., Cai, Z., & Zhang, T. (2025). Extreme winter environment dominates gut microbiota and metabolome of white-lipped deer. Microbiological Research, 297, 128182. 10.1016/j.micres.2025.128182

Li, D., & Gaquerel, E. (2021). Next-Generation Mass Spectrometry Metabolomics Revives the Functional Analysis of Plant Metabolic Diversity. Annual Review of Plant Biology, 72, 867–891. 10.1146/annurev-arplant-071720-114836

Li, L., Tian, Z., Chen, J., Tan, Z., Zhang, Y., Zhao, H., Wu, X., Yao, X., Wen, W., Chen, W., & Guo, L. (2023). Characterization of novel loci controlling seed oil content in Brassica napus by marker metabolite-based multi-omics analysis. Genome Biology, 24(1), 141. 10.1186/s13059-023-02984-z

Liska, O., Boross, G., Rocabert, C., Szappanos, B., Tengölics, R., & Papp, B. (2023). Principles of metabolome conservation in animals. Proceedings of the National Academy of Sciences, 120(35), e2302147120. 10.1073/pnas.2302147120

Liu, Y., Watanabe, M., Yasukawa, S., Kawamura, Y., Aneklaphakij, C., Fernie, A. R., & Tohge, T. (2021). Cross-Species Metabolic Profiling of Floral Specialized Metabolism Facilitates Understanding of Evolutional Aspects of Metabolism Among Brassicaceae Species. Frontiers in Plant Science, 12. 10.3389/fpls.2021.640141

Macias, S., Yilmaz, A., Kirma, J., Moore, S. E., Woodside, J. V., Graham, S. F., & Green, B. D. (2023). Non-targeted LC–MS/MS metabolomic profiling of human plasma uncovers a novel Mediterranean diet biomarker panel. Metabolomics, 20(1), 3.

Matthews, B., Marchinko, K.B., Bolnick, D.I. and Mazumder, A., 2010. Specialization of trophic position and habitat use by sticklebacks in an adaptive radiation. Ecology, 91(4), pp.1025–1034.

Martino, C., Morton, J.T., Marotz, C.A., Thompson, L.R., Tripathi, A., Knight, R., Zengler, K,. 2019. A Novel Sparse Compositional Technique Reveals Microbial Perturbations. mSystems4:10.1128/msystems.00016-19. 10.1128/msystems.00016-19

McPhail, J.D. 1984. Ecology and evolution of sympatric sticklebacks (Gasterosteus): morphological and genetic evidence for a species pair in Enos Lake, British Columbia. Canadian Journal of Zoology, 62(7): 1402–1408. doi:10.1139/z84-201.

Nomoto, H., Fernández-Conradi, P., Kjelsberg, N., Defossez, E., Münzbergová, Z., Glauser, G., & Rasmann, S. (2025). Elevation drives intraspecific metabolomic differentiation in natural and experimental populations. Plant Biology, 27(5), 773–784. 10.1111/plb.70025

Oksanen, J., Simpson, G. L., Blanchet, F. G., Kindt, R., Legendre, P., Minchin, P. R., O’Hara, R. B., Solymos, P., Stevens, M. H. H., Szoecs, E., Wagner, H., Barbour, M., Bedward, M., Bengtsson, H., Bolker, B., Borcard, D., Borman, T., Carvalho, G., Chirico, M., … Weedon, J. (2026). vegan: Community Ecology Package (Version 2.7-5) [Computer software]. https://cran.r-project.org/web/packages/vegan/index.html

Oliver, S. G., Winson, M. K., Kell, D. B., & Baganz, F. (1998). Systematic functional analysis of the yeast genome. Trends in Biotechnology, 16(9), 373–378. 10.1016/S0167-7799(98)01214-1

Parthasarathy, A., Cross, P. J., Dobson, R. C. J., Adams, L. E., Savka, M. A., & Hudson, A. O. (2018). A Three-Ring Circus: Metabolism of the Three Proteogenic Aromatic Amino Acids and Their Role in the Health of Plants and Animals. Frontiers in Molecular Biosciences, 5, 29. 10.3389/fmolb.2018.00029

Patti, G. J., Yanes, O., & Siuzdak, G. (2012). Metabolomics: The apogee of the omics trilogy. Nature Reviews Molecular Cell Biology, 13(4), 263–269. 10.1038/nrm3314

Pullman, B., Batsoyol, N., Wang, M., Swanson, S. and Bandeira, N. (2021) Real-time modification-tolerant matching of MS/MS spectra at the repository scale. Presented at: ASMS Annual Conference, MP159.

Quinn, R. A., Melnik, A. V., Vrbanac, A., Fu, T., Patras, K. A., Christy, M. P., Bodai, Z., Belda-Ferre, P., Tripathi, A., Chung, L. K., Downes, M., Welch, R. D., Quinn, M., Humphrey, G., Panitchpakdi, M., Weldon, K. C., Aksenov, A., da Silva, R., Avila-Pacheco, J., … Dorrestein, P. C. (2020). Global chemical effects of the microbiome include new bile-acid conjugations. Nature, 579(7797), 123–129. 10.1038/s41586-020-2047-9

Ramanathan, C., Goris, L., Mishra, A., Lihavainen-Bag, J., Pawlowski, K., Albrectsen, B. R., & Tack, A. J. M. (2026). The Effects of Acorn Origin, Environmental Microbiomes and Local Adaptation on the Leaf Metabolome. Journal of Chemical Ecology, 52(1), 18. 10.1007/s10886-026-01692-9

Rao, M. J., Wang, H., Lei, H., Zhang, H., Duan, X., Bao, L., Yang, C., Han, D., Zhang, Y., Yang, S., & Duan, M. (2024). LC-MS/MS-based metabolomic study provides insights into altitude-dependent variations in flavonoid profiles of strawberries. Frontiers in Plant Science, 15, 1527212. 10.3389/fpls.2024.1527212

Rattray, N. J. W., Deziel, N. C., Wallach, J. D., Khan, S. A., Vasiliou, V., Ioannidis, J. P. A., & Johnson, C. H. (2018). Beyond genomics: Understanding exposotypes through metabolomics. Human Genomics, 12(1), 4. 10.1186/s40246-018-0134-x

Reid, K., Bell, M. A., & Veeramah, K. R. (2021). Threespine Stickleback: A Model System For Evolutionary Genomics. Annual Review of Genomics and Human Genetics, 22, 357–383. 10.1146/annurev-genom-111720-081402

Ridlon, J. M., Wolf, P. G., & Gaskins, H. R. (2016). Taurocholic acid metabolism by gut microbes and colon cancer. Gut Microbes, 7(3), 201–215. 10.1080/19490976.2016.1150414

Ritchie, M. E., Phipson, B., Wu, D., Hu, Y., Law, C. W., Shi, W., & Smyth, G. K. (2015). Limma powers differential expression analyses for RNA-sequencing and microarray studies. Nucleic Acids Research, 43(7), e47. 10.1093/nar/gkv007

Rohart, F., Gautier, B., Singh, A., & Cao, K.-A. L. (2017). mixOmics: An R package for ‘omics feature selection and multiple data integration. PLOS Computational Biology, 13(11), e1005752. 10.1371/journal.pcbi.1005752

Ryan, D., & Robards, K. (2006). Metabolomics: The greatest omics of them all? Analytical Chemistry, 78(23), 7954–7958. 10.1021/ac0614341

Sarkar, J., Singh, R., & Chandel, S. (2025). Understanding LC/MS-Based Metabolomics: A Detailed Reference for Natural Product Analysis. PROTEOMICS – Clinical Applications, 19(1), e202400048. 10.1002/prca.202400048

Schluter, D., & McPhail, J. D. (1992). Ecological Character Displacement and Speciation in Sticklebacks. The American Naturalist, 140(1), 85–108. 10.1086/285404

Schmid, R., Heuckeroth, S., Korf, A., Smirnov, A., Myers, O., Dyrlund, T. S., Bushuiev, R., Murray, K. J., Hoffmann, N., Lu, M., Sarvepalli, A., Zhang, Z., Fleischauer, M., Dührkop, K., Wesner, M., Hoogstra, S. J., Rudt, E., Mokshyna, O., Brungs, C., … Pluskal, T. (2023). Integrative analysis of multimodal mass spectrometry data in MZmine 3. Nature Biotechnology, 41(4), 447–449. 10.1038/s41587-023-01690-2

Schrieber, K., Glüsing, S., Peters, L., Eichert, B., Althoff, M., Schwarz, K., Erfmeier, A., & Demetrowitsch, T. (2023). Population divergence in heat and drought responses of a coastal plant: From metabolic phenotypes to plant morphology and growth. Journal of Experimental Botany, 74(15), 4559–4578. 10.1093/jxb/erad147

Schymanski, E. L., Jeon, J., Gulde, R., Fenner, K., Ruff, M., Singer, H. P., & Hollender, J. (2014). Identifying Small Molecules via High Resolution Mass Spectrometry: Communicating Confidence. Environmental Science & Technology, 48(4), 2097–2098. 10.1021/es5002105

Sedio, B. E. (2017). Recent breakthroughs in metabolomics promise to reveal the cryptic chemical traits that mediate plant community composition, character evolution and lineage diversification. New Phytologist, 214(3), 952–958. 10.1111/nph.14438

Sedio, B. E., Parker, J. D., McMahon, S. M., & Wright, S. J. (2018). Comparative foliar metabolomics of a tropical and a temperate forest community. Ecology, 99(12), 2647–2653. 10.1002/ecy.2533

Shannon, P., Markiel, A., Ozier, O., Baliga, N. S., Wang, J. T., Ramage, D., Amin, N., Schwikowski, B., & Ideker, T. (2003). Cytoscape: A Software Environment for Integrated Models of Biomolecular Interaction Networks. Genome Research, 13(11), 2498–2504. 10.1101/gr.1239303

Shende, V. V., Bauman, K. D., & Moore, B. S. (2024). The shikimate pathway: Gateway to metabolic diversity. Natural Product Reports, 41(4), 604–648. 10.1039/D3NP00037K

Slowikowski K (2026). ggrepel: Automatically Position Non-Overlapping Text Labels with ’ggplot2’. R package version 0.9.8, https://ggrepel.slowkow.com/.

Sparagon, W. J., Gentry, E. C., Minich, J. J., Vollbrecht, L., Laurens, L. M. L., Allen, E. E., Sims, N. A., Dorrestein, P. C., Kelly, L. W., & Nelson, C. E. (2022). Fine scale transitions of the microbiota and metabolome along the gastrointestinal tract of herbivorous fishes. Animal Microbiome, 4(1), 33. 10.1186/s42523-022-00182-z

Strickland, K., Matthews, B., Jónsson, Z. O., Kristjánsson, B. K., Phillips, J. S., Einarsson, Á., & Räsänen, K. (2024). Microevolutionary change in wild stickleback: Using integrative time-series data to infer responses to selection. Proceedings of the National Academy of Sciences, 121(37), e2410324121. 10.1073/pnas.2410324121

Stuart, Y. E., Veen, T., Weber, J. N., Hanson, D., Ravinet, M., Lohman, B. K., Thompson, C. J., Tasneem, T., Doggett, A., Izen, R., Ahmed, N., Barrett, R. D. H., Hendry, A. P., Peichel, C. L., & Bolnick, D. I. (2017). Contrasting effects of environment and genetics generate a continuum of parallel evolution. Nature Ecology & Evolution, 1(6), 0158. 10.1038/s41559-017-0158

Sumner, L. W., Amberg, A., Barrett, D., Beale, M. H., Beger, R., Daykin, C. A., Fan, T. W.-M., Fiehn, O., Goodacre, R., Griffin, J. L., Hankemeier, T., Hardy, N., Harnly, J., Higashi, R., Kopka, J., Lane, A. N., Lindon, J. C., Marriott, P., Nicholls, A. W., … Viant, M. R. (2007). Proposed minimum reporting standards for chemical analysis. Metabolomics, 3(3), 211–221. 10.1007/s11306-007-0082-2

Tahir, U. A., Katz, D. H., Avila-Pachecho, J., Bick, A. G., Pampana, A., Robbins, J. M., Yu, Z., Chen, Z.-Z., Benson, M. D., Cruz, D. E., Ngo, D., Deng, S., Shi, X., Zheng, S., Eisman, A. S., Farrell, L., Hall, M. E., Correa, A., Tracy, R. P., … Gerszten, R. E. (2022). Whole Genome Association Study of the Plasma Metabolome Identifies Metabolites Linked to Cardiometabolic Disease in Black Individuals. Nature Communications, 13(1), 4923. 10.1038/s41467-022-32275-3

Tian, J., Zhu, X., Wu, H., Wang, Y., & Hu, X. (2023). Serum metabolic profile and metabolome genome-wide association study in chicken. Journal of Animal Science and Biotechnology, 14(1), 69. 10.1186/s40104-023-00868-7

Tudor, E. P., Lewandrowski, W., Krauss, S., & Veneklaas, E. J. (2024). Local adaptation to climate inferred from intraspecific variation in plant functional traits along a latitudinal gradient. Conservation Physiology, 12(1), coae018. 10.1093/conphys/coae018

Wang, J., Feng, Y., Xu, S., Tenzin, N., Han, H., Gong, D., Liu, F., Sun, Y., & Liu, S. (2025). Non-targeted LC-MS metabolomics reveals serum metabolites for high-altitude adaptation in Tibetan donkeys. Scientific Reports, 15(1), 46. 10.1038/s41598-024-83544-8

Wang, M., Carver, J. J., Phelan, V. V., Sanchez, L. M., Garg, N., Peng, Y., Nguyen, D. D., Watrous, J., Kapono, C. A., Luzzatto-Knaan, T., Porto, C., Bouslimani, A., Melnik, A. V., Meehan, M. J., Liu, W.-T., Crüsemann, M., Boudreau, P. D., Esquenazi, E., Sandoval-Calderón, M., … Bandeira, N. (2016). Sharing and community curation of mass spectrometry data with Global Natural Products Social Molecular Networking. Nature Biotechnology, 34(8), 828–837. 10.1038/nbt.3597

Wang, M., Wang, J., Carver, J., Pullman, B. S., Cha, S. W., & Bandeira, N. (2018). Assembling the Community-Scale Discoverable Human Proteome. Cell Systems, 7(4), 412–421.e5. 10.1016/j.cels.2018.08.004

Wieder, C., Frainay, C., Poupin, N., Rodríguez-Mier, P., Vinson, F., Cooke, J., Lai, R. P., Bundy, J. G., Jourdan, F., & Ebbels, T. (2021). Pathway analysis in metabolomics: Recommendations for the use of over-representation analysis. PLoS Computational Biology, 17(9), e1009105. 10.1371/journal.pcbi.1009105

Willacker, J. J., von Hippel, F. A., Wilton, P. R., & Walton, K. M. (2010). Classification of threespine stickleback along the benthic-limnetic axis. Biological Journal of the Linnean Society. Linnean Society of London, 101(3), 595–608. 10.1111/j.1095-8312.2010.01531.x

Wolters, F. C., Woldu, T., Schranz, E. M., Medema, M. H., Bouwmeester, K., & Hooft, J. J. J. van der. (2026). Metabolic fingerprinting of 17 Brassicaceae species across three tissues (p. 2026.04.17.719198). bioRxiv. 10.64898/2026.04.17.719198

Wu, X.-X., Ban, W.-L., Wu, L.-J., Qi, W.-J., Borhani, M., He, X.-Y., Liu, X.-L., Liu, M.-Y., & Ding, J. (2024). Identification of serum biomarkers for cystic echinococcosis in sheep through untargeted metabolomic analysis using LC-MS/MS technology. Parasites & Vectors, 17(1), 547. 10.1186/s13071-024-06599-6

Xie, H., Hu, J., Zhao, X., Chen, J., Yue, X., Zhou, C., Navarro-Muñoz, J. C., Jiang, J., Tang, X., Zhao, F., Hatmaker, E. A., Rokas, A., Barber, A. E., Drott, M. T., Keller, N. P., Zhang, Q., van der Hooft, J. J. J., Medema, M. H., & Li, P. (2026a). Large-scale multi-omics profiling reveals environmental and evolutionary drivers of fungal phylogeographic and metabolic diversity. Nature Communications, 17(1), 4121. 10.1038/s41467-026-70721-8

Xie, H., Hu, J., Zhao, X., Chen, J., Yue, X., Zhou, C., Navarro-Muñoz, J. C., Jiang, J., Tang, X., Zhao, F., Hatmaker, E. A., Rokas, A., Barber, A. E., Drott, M. T., Keller, N. P., Zhang, Q., van der Hooft, J. J. J., Medema, M. H., & Li, P. (2026b). Large-scale multi-omics profiling reveals environmental and evolutionary drivers of fungal phylogeographic and metabolic diversity. Nature Communications, 17(1), 4121. 10.1038/s41467-026-70721-8

Xu, H., Zhang, Q., Kim, S.-K., Liao, Z., Wei, Y., Sun, B., Jia, L., Chi, S., & Liang, M. (2020). Dietary taurine stimulates the hepatic biosynthesis of both bile acids and cholesterol in the marine teleost, tiger puffer (Takifugu rubripes). The British Journal of Nutrition, 123(12), 1345–1356. 10.1017/S0007114520000161

Xu, J., Xie, S., Chi, S., Zhang, S., Cao, J., & Tan, B. (2022). Protective effects of taurocholic acid on excessive hepatic lipid accumulation via regulation of bile acid metabolism in grouper. Food & Function, 13(5), 3050–3062. 10.1039/D1FO04085E

Xu, W., Meng, Z., Deng, J., Sun, X., Liu, T., Tang, Y., Zhang, Z., Liu, Y., & Zhu, W. (2022). Metabonomic identification of serum biomarkers related to heat stress tolerance of sheep. Animal Science Journal = Nihon Chikusan Gakkaiho, 93(1), e13792. 10.1111/asj.13792

Yasaka, T. M., Kim, C. K., Meadows, V., & Monga, S. P. (2026). Zonation, Zonation, Zonation: The Real Estate of the Liver. Annual Review of Pathology, 21(1), 185–212. 10.1146/annurev-pathmechdis-042624-091820

Zhang, M., You, M., Ma, N., & Lv, J. (2024). Advance in the application of metabolomics technology in poultry. Frontiers in Veterinary Science, 11. 10.3389/fvets.2024.1501630

Zuffa, S., Charron-Lamoureux, V., Brennan, C., Ambre, M., Knight, R., & Dorrestein, P. C. (2025). Human Untargeted Metabolomics in High-Throughput Gut Microbiome Research: Ethanol vs Methanol. Analytical Chemistry, 97(9), 4945–4953. 10.1021/acs.analchem.4c05142

